# LD Score Regression Distinguishes Confounding from Polygenicity in Genome-Wide Association Studies

**DOI:** 10.1101/002931

**Authors:** Brendan K. Bulik-Sullivan, Po-Ru Loh, Hilary Finucane, Stephan Ripke, Jian Yang, Schizophrenia Working Group of the Psychiatric Genomics Consortium, Nick Patterson, Mark J. Daly, Alkes L. Price, Benjamin M. Neale

## Abstract

Both polygenicity^1,2^ (*i.e.* many small genetic effects) and confounding biases, such as cryptic relatedness and population stratification^3^, can yield inflated distributions of test statistics in genome-wide association studies (GWAS). However, current methods cannot distinguish between inflation from bias and true signal from polygenicity. We have developed an approach that quantifies the contributions of each by examining the relationship between test statistics and linkage disequilibrium (LD). We term this approach LD Score regression. LD Score regression provides an upper bound on the contribution of confounding bias to the observed inflation in test statistics and can be used to estimate a more powerful correction factor than genomic control^4–14^. We find strong evidence that polygenicity accounts for the majority of test statistic inflation in many GWAS of large sample size.

---

Variants in LD with a causal variant show elevated test statistics in association analysis proportional to the LD (measured by *r^2^)* with the causal variant^1,15,16^. The more genetic variation an index variant tags, the higher the probability that this index variant will tag a causal variant. Precisely, if we assume that effect sizes are drawn independently from distributions with variance proportional to 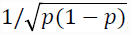, where *p* is minor allele frequency (MAF), then the expected *χ*^2^-statistic of variant *j* is

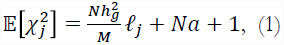

where *N* is sample size;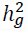 is the squared correlation (in the population) between phenotype and the best linear predictor that can be constructed from genotyped SNPs; *M* is the number of variants; 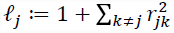 is the LD Score of variant *j*, which measures the amount of genetic variation tagged by *j* and *a* measures the contribution of confounding bias. The same relationship holds for ascertained studies of binary phenotypes if we replace 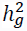 with observed scale heritability (Supplementary Note). In contrast, inflation from cryptic relatedness within or between cohorts^4,17,18^ or population stratification from genetic drift will not correlate with LD Score. Consequently, if we regress *χ*^2^-statistics from GWAS against LD Score (LD Score regression), the intercept from this regression will estimate the mean contribution of confounding bias to the inflation in the test statistics, and a statistically significant positive slope is evidence of real polygenic signal (Supplementary Note).

We briefly summarize LD Score estimation and regression here, with a more thorough description in the Online Methods. We estimated LD Scores from the European ancestry samples in the 1000 Genomes Project ^19^ (EUR) using an unbiased estimator ^21^ of *r^2^* with 1 centiMorgan (cM) windows, singletons excluded (MAF > 0.13%) and no *r^2^* cutoff. Standard errors were estimated by jackknifing over samples, and we used these standard errors to correct for attenuation bias (*i.e.,* the downward bias in the magnitude of the regression slope that results when the regressor is measured noisily) in LD Score regression. In addition, we excluded variants with EUR MAF < 1% from all regressions because the LD Score standard errors for these variants were very high (note: these variants were included in LD Score estimation. In general, it is important to distinguish between the set of variants included in the sum of *r*^*2*^, *s* and the set of variants included in the regression, since the latter will often be a subset of the former).

Mismatch between LD Scores in a reference population used for LD Score estimation and a target population used for GWAS can bias the LD Score regression in two ways. First, if LD Scores in the reference population are noisy approximations to LD Scores in the target population, then our standard errors will underestimate the standard error of 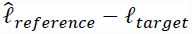, which will bias the LD Score regression slope downwards and the intercept upwards. Second, if LD Scores in the reference population are systematically higher or lower than in the target population, this can bias the LD Score regression intercept downwards or upwards, respectively. In order to quantify the extent of intra-European differences in LD Score, we estimated LD Scores using each of the 1000 Genomes EUR subpopulations (Utah Residents with Northern and Western European Ancestry (CEU), British in England and Scotland (GBR), Toscani in Italia (TSI) and Finnish in Finland (FIN)) separately. The LD Scores from all four subpopulations were highly correlated, but mean LD Score increased with latitude, consistent with the observation that Southern European populations have gone through less severe bottlenecks than Northern European populations^22^. For example, the mean LD Score for FIN was 7% larger and the mean LD Score for TSI was 8% smaller than for EUR. Nevertheless, the EUR reference panel is appropriate for studies in outbred populations of predominantly northern European ancestry, such as European American or UK populations. For other populations, a different reference panel should be used.

We performed a variety of simulations to verify the relationship between linkage disequilibrium and *χ*^2^-statistics under scenarios with population stratification, cryptic relatedness and polygenic architecture. To simulate population stratification on a continental scale, we obtained un-imputed genotypes from Psychiatric Genomics Consortium (PGC) controls from seven European samples, all genotyped on the same array (Supplementary Table 1). For each pair of samples, we assigned case/control status based on sample membership, then computed association statistics using PLINK^23^ (Online Methods). To model population stratification on a national scale, we computed the top three principal components within each cohort, then computed association statistics using each of these principal components as phenotypes.

**Figure 1.**
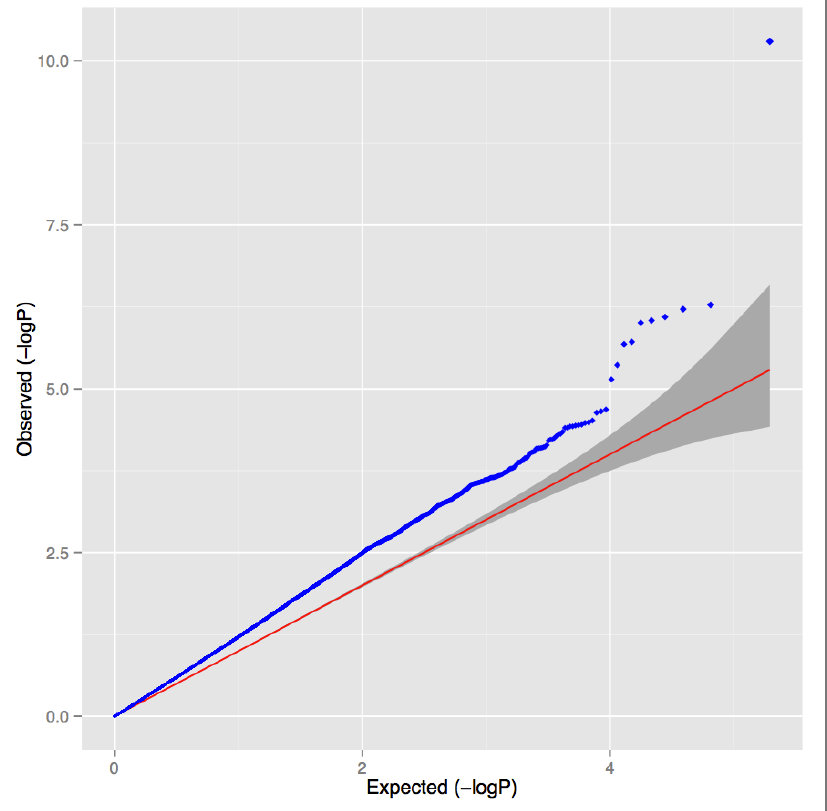
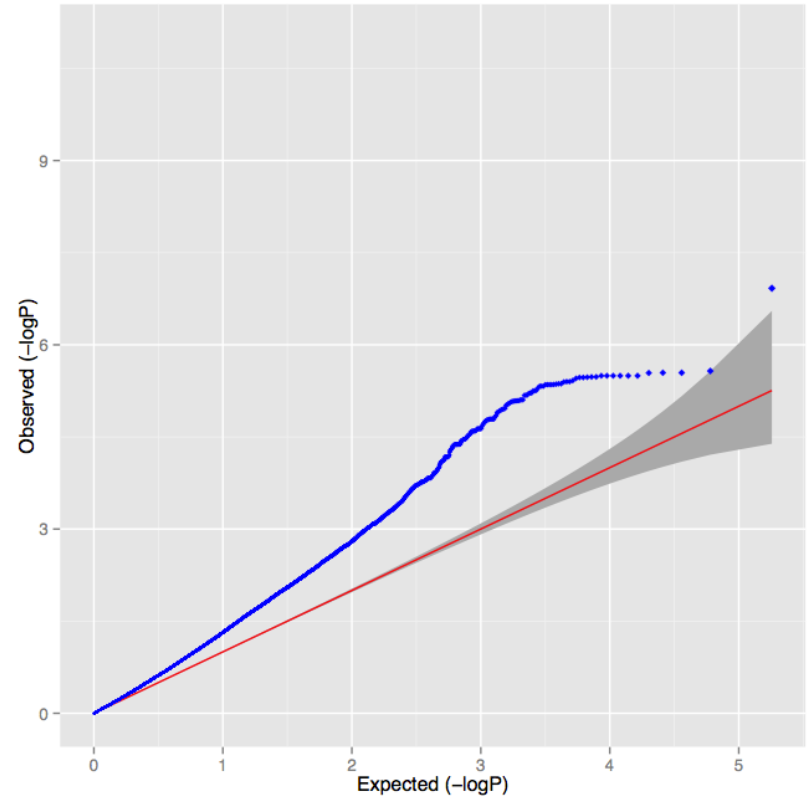
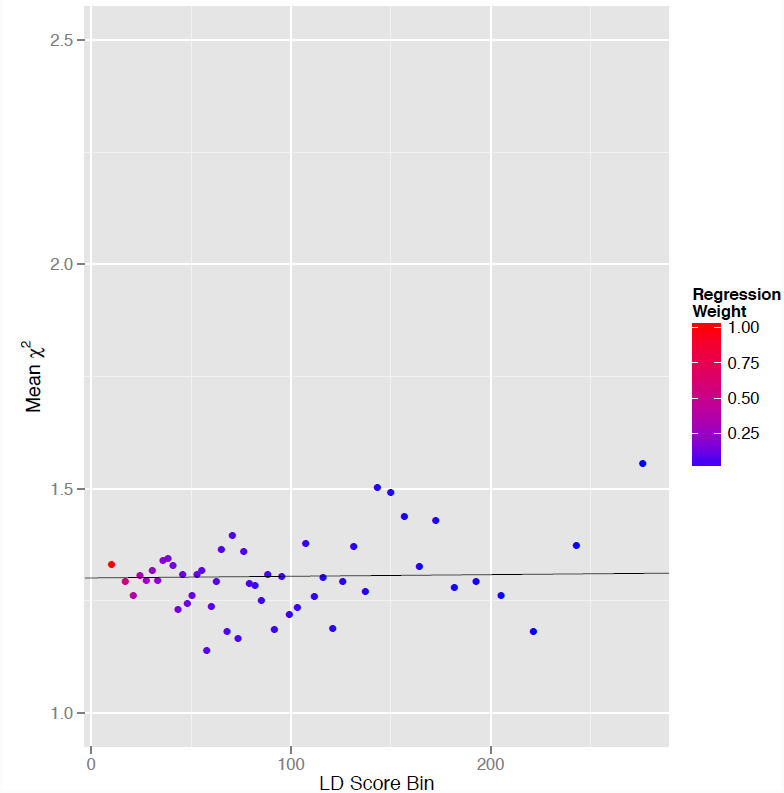
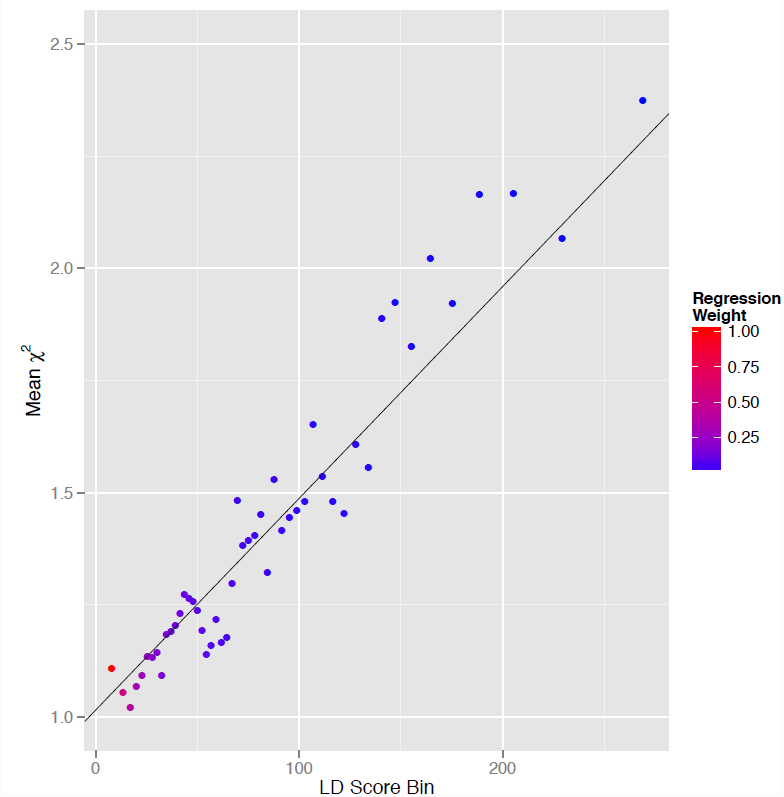
Results from selected simulations. **(a)** QQ plot with population stratification (*λ_GC_* 1.32, LD Score regression intercept = 1.30). **(b)** QQ plot with polygenic genetic architecture with 0.1% of SNPs causal (*λ_GC_* = 1.32, LD Score regression intercept = 1.02) **(c)** LD Score plot with population stratification. Each point represents an LD Score quantile, where the *x*-coordinate of the point is the mean LD Score of variants in that quantile and the *y*-coordinate is the mean *χ*^2^ of variants in that quantile. Colors correspond to regression weights, with red indicating large weight. The black line is the LD Score regression line, block jackknife *p*-value = 0.45 **(d)** As in panel c but LD Score plot with polygenic genetic architecture, block jackknife *p*-value = 1.6 × 10^−69^.

To model a polygenic quantitative trait, we assigned effect sizes drawn from 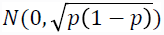 to varying numbers of causal variants in an approximately unstructured sample of 1000 Swedes. Quantile-quantile (QQ) plots from simulations with population stratification and polygenicity show indistinguishable patterns of inflation (Fig. 1a,b). However, the LD Score regression intercept was consistently near the mean *χ^2^* in simulations with pure stratification (Fig. 1c, Supplementary Tables 2-3) and near 1 in simulations with polygenicity (Fig. 1d, Supplementary Figures 1-5), demonstrating that LD Score regression can distinguish between the two. We note that if there are few causal variants, then the LD Score regression becomes unstable (Supplementary Figure 6).

To simulate a more realistic scenario where both polygenicity and bias contribute simultaneously to test statistic inflation, we obtained genotypes from approximately 22,000 individuals from throughout Europe from the Wellcome Trust Case-Control Consortium 2^24^. We simulated polygenic phenotypes with causal SNPs drawn from the first halves of chromosomes, leaving all SNPs on the second halves of chromosomes null. In addition, we included an environmental stratification component aligned with the first principal component of the genotype data, representing Northern vs. Southern European ancestry. In this setup, the mean *χ*^2^ among SNPs on the second halves of chromosomes measures the average contribution of stratification. We performed similar simulations with cryptic relatedness using data from the Framingham Heart Study^25^, which includes close relatives. Results from these simulations are summarized in Supplementary Table 4. In all simulation replicates the LD Score regression intercept was approximately equal to the mean *χ*^2^ among null SNPs, which demonstrates that LD Score regression can partition the inflation in test statistics even in the presence of both bias and polygenicity. Finally, we modeled studies of a polygenic binary phenotype with case ascertainment using a simulated genotypes and a liability threshold model, and verified that LD Score regression is not noticeably biased by ascertainment (Supplementary Table 5).

A potential limitation of LD Score regression is that effect sizes may be correlated with LD Score. For instance, this could occur if variance explained is correlated with MAF, because MAF is correlated with LD Score. To quantify the magnitude of the bias that this would introduce, we simulated a variety of frequency-dependent genetic architectures (Online Methods). Under genetic architectures where mean variance explained was highest among rare variants, the LD Score regression intercept was biased upwards, and vice versa when mean variance explained was highest for common variants. The magnitude of this effect was small for reasonable genetic architectures; for instance, under a genetic architecture with 0.25% of SNPs causal and variance explained proportional to 1/*p*(1 − *p*), the mean inflation in the LD Score regression intercept across 10 simulations was only 11% of the mean *χ^2^*, and under a genetic architecture with variance explained proportional to *p*(1 − *p*), the mean LD Score regression intercept was 0.994 (Supplementary Figure 6, Supplementary Table 6). However, LD Score regression is not effective for extreme genetic architectures: if all causal variants are rare (MAF < 1%), then LD Score regression will often generate a negative slope, and the intercept will be exceed the mean *χ^2^* (Supplementary Figure 7).

**Table 1.** LD Score regression results for all studies analyzed that either did not apply meta-analysis level GC correction or listed *λ_GC_* in the relevant publication. The column labeled “GC” indicates how many rounds of GC correction were performed. For GWAS that applied meta-analysis level GC correction and listed *λ_GC_*, we re-inflated all test statistics by the meta-analysis level *λ_GC_*. Results after double GC correction are displayed in Supplementary Table 8. Standard errors and *p*-values are obtained via a block jackknife over blocks of ~2000 adjacent SNPs, which provides a robust estimate of standard error in the presence of correlated, heteroskedastic error terms. The column labeled “Type” indicates whether the study was a mega- (raw genotypes shared between studies) or meta-analysis (only summary statistics shared between all contributing studies).

| Phenotype | Mean $\chi^2$ | $\lambda_{GC}$ | Intercept | Intercept SE | Slope | Slope SE | Type | GC | P-Value | Ref |
| --- | --- | --- | --- | --- | --- | --- | --- | --- | --- | --- |
| Inflammatory bowel disease | 1.247 | 1.164 | 1.086 | 0.006435 | 0.001545 | 8.47E-05 | mega | 0 | 1.34E-74 | 40 |
| Ulcerative Colitis | 1.174 | 1.128 | 1.071 | 0.006242 | 0.0009801 | 7.86E-05 | mega | 0 | 5.47E-36 | 40 |
| Crohn's | 1.185 | 1.122 | 1.037 | 0.005237 | 0.001415 | 7.23E-05 | mega | 0 | 1.02E-85 | 40 |
| Schizophrenia | 1.613 | 1.484 | 1.066 | 0.00563 | 0.005209 | 8.01E-05 | mega | 0 | <1E-300 | 26 |
| ADHD | 1.033 | 1.033 | 1.006 | 0.005363 | 0.0002144 | 6.65E-05 | mega | 0 | 6.31E-04 | 41 |
| Bipolar | 1.154 | 1.135 | 1.028 | 0.005745 | 0.001220 | 7.27E-05 | mega | 0 | 1.36E-63 | 42 |
| PGC Cross-Disorder | 1.205 | 1.187 | 1.013 | 0.005779 | 0.001869 | 7.56E-05 | mega | 0 | 2.78E-135 | 43 |
| Major Depression | 1.063 | 1.063 | 1.005 | 0.005226 | 0.0005718 | 6.45E-05 | mega | 0 | 3.89E-19 | 44 |
| Rheumatoid Arthritis | 1.063 | 1.033 | 0.9798 | 0.0007542 | 0.0007252 | 6.50E-05 | mega | 2 | 2.20E-31 | 8 |
| Coronary Artery Disease | 1.125 | 1.096 | 1.027 | 0.005270 | 0.0009507 | 6.58E-05 | meta | 1 | 1.22E-47 | 11 |
| Type-2 Diabetes | 1.116 | 1.097 | 1.021 | 0.005614 | 0.0008802 | 6.95E-05 | meta | 1 | 4.68E-37 | 45 |
| BMI-Adj. Fasting Insulin | 1.088 | 1.072 | 1.009 | 0.005208 | 0.0007578 | 6.59E-05 | meta | 1 | 6.87E-31 | 12 |
| Fasting Insulin | 1.079 | 1.067 | 1.019 | 0.005220 | 0.0005517 | 6.54E-05 | meta | 1 | 1.65E-17 | 12 |
| College | 1.207 | 1.180 | 1.041 | 0.005484 | 0.001545 | 6.50E-05 | meta | 1 | 3.17E-125 | 13 |
| Years of Education | 1.220 | 1.188 | 1.040 | 0.005666 | 0.001660 | 6.65E-05 | meta | 1 | 8.57E-138 | 13 |
| Cigarettes Per Day | 1.047 | 1.047 | 0.9968 | 0.005554 | 0.0005349 | 7.02E-05 | meta | 1 | 1.29E-14 | 9 |
| Ever Smoked | 1.097 | 1.083 | 0.9987 | 0.005201 | 0.0009647 | 6.48E-05 | meta | 1 | 1.70E-50 | 9 |
| Former Smoker | 1.050 | 1.048 | 0.9953 | 0.005249 | 0.0005366 | 6.57E-05 | meta | 1 | 1.65E-16 | 9 |
| Age-Onset (Smoking) | 1.025 | 1.030 | 0.9966 | 0.005212 | 0.0002668 | 6.56E-05 | meta | 1 | 2.39E-05 | 9 |
| FN-BMD | 1.163 | 1.109 | 0.9934 | 0.001613 | 0.001450 | 7.15E-05 | meta | 2 | 5.49E-113 | 14 |
| LS-BMD | 1.174 | 1.112 | 1.022 | 0.001444 | 0.001281 | 7.27E-05 | meta | 2 | 6.04E-88 | 14 |
| Waist-Hip Ratio | 1.417 | 1.330 | 1.040 | 0.003633 | 0.002756 | 8.46E-05 | meta | 2 | <1E-300 | 5 |
| Height | 1.802 | 1.478 | 1.145 | 0.006229 | 0.004387 | 1.20E-04 | meta | 2 | <1E-300 | 6 |
| Body-Mass Index | 1.130 | 1.090 | 1.025 | 0.0009696 | 0.0008895 | 6.35E-05 | meta | 2 | 5.24E-53 | 7 |

Next, we applied LD Score regression to summary statistics from GWAS representing more than 30 different phenotypes ^4–6,16–29^ (Online Methods, Table 1, Supplementary Tables 8,10,11). For all studies, the slope of the LD Score regression was significantly greater than 0, and the LD Score regression intercept was significantly less than *λ_GC_* (mean difference 0.11), suggesting that polygenicity significantly contributes to the increase in mean *χ^2^* and confirming that correcting test statistics by dividing by *λ_GC_* is unnecessarily conservative. As an example, Figure 2 displays the LD Score regression for the most recent schizophrenia GWAS, restricted to ~70,000 European individuals^26^. The low intercept of 1.066 and highly significant slope (*p*<10^−300^) indicate at most a small contribution of bias, and that the mean *χ^2^* of 1.613 results mostly from polygenicity.

**Figure 2.**
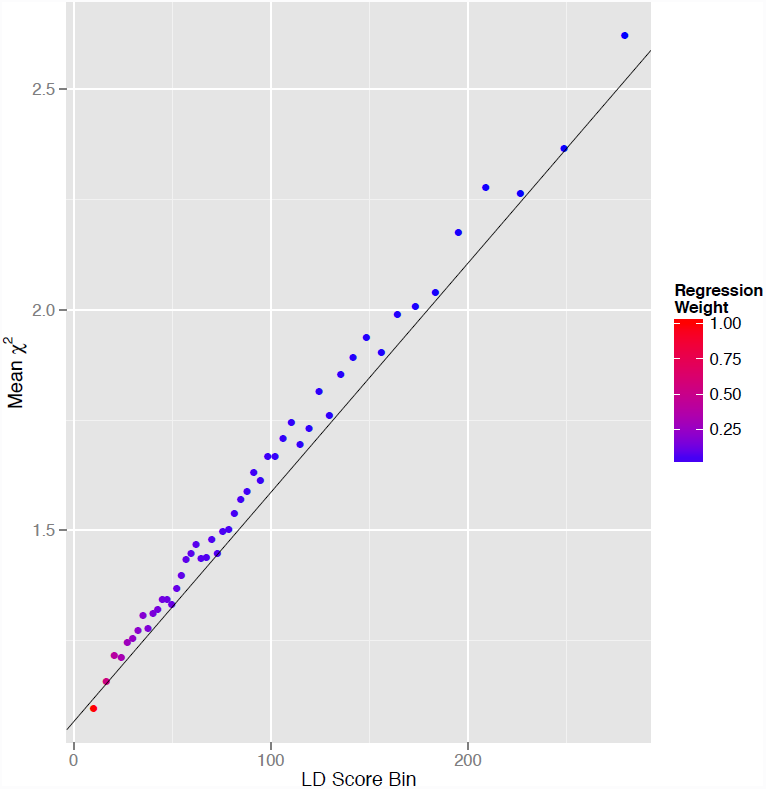
LD Score regression plot for the current schizophrenia meta-analysis^26^. Each point represents an LD Score quantile, where the ***x***-coordinate of the point is the mean LD Score of variants in that quantile and the *y*-coordinate is the mean *χ^2^* of variants in that quantile. Colors correspond to regression weights, with red indicating large weight. The black line is the LD Score regression line, block jackknife *p*-value < 10^−300^.

Finally, we discuss application of the LD Score regression intercept as a correction factor for GWAS meta-analyses. Where possible, it is preferable to obtain all genotype data and correct for confounding biases directly ^27–31^; however, if only summary data are available, or if a conservative correction is desired, we propose that the LD Score regression intercept provides a more robust upper bound on the extent of inflation from confounding bias than *λ_GC_* (or intergenic *λ_GC_*, Supplementary Table 9). Since *λ_GC_* increases with sample size even in the absence of confounding bias^1^, the gain in power obtained by correcting test statistics with the LD Score regression intercept instead of *λ_GC_* will become even more substantial for larger GWAS. Extending this method to non-European populations such as East Asians or West Africans is straightforward given appropriate reference panels, but extension to admixed populations is the subject of future research.

In conclusion, we have developed LD Score regression, a method to distinguish between inflated test statistics from confounding bias and polygenicity. Application of LD Score regression to over 30 complex traits confirms that polygenicity accounts for the majority of test statistic inflation in GWAS results and this approach can be used to generate a correction factor for GWAS that is more powerful than *λ*_GC_, especially at large sample sizes. We have made an R script for performing LD Score regression and a database of LD Scores suitable for European-ancestry samples available for download (URLs). Research in progress aims to apply this method to estimation of heritability and genetic correlation, as well as to the calibration of mixed model statistics.

## ONLINE METHODS

### Estimation of LD Score and Standard Error

We estimated European LD Scores from 378 phased European individuals (excluding one individual from a pair of cousins) from the 1000 Genomes Project reference panel using the –ld-mean-rsq option implemented in the GCTA^20^ software package (with flags –ld-mean-rsq –ld-rsq-cutoff 0 –maf 0.00001; we implemented a 1centiMorgan (cM) window using the –ld-wind flag and modified .bim files with physical coordinates replaced with genetic coordinates as described in the next paragraph). The primary rationale for using a sequenced reference panel containing several hundred individuals for LD Score estimation rather than a genotyped GWAS control panel with several thousand individuals was that even after imputing off-chip genotypes, the variants available from a genotyping array only account for a subset of all variants. Using only a subset of all variants for estimating LD Score produces estimates that are biased downwards.

We used a window of radius 1cM around the index variant for the sum of *r^2^*’s (using the genetic map and phased genotypes from the IMPUTE2 website, see URLs), no *r^2^* cutoff, and excluded singletons (MAF < 0.13%). The standard estimator of the Pearson correlation coefficient has upward bias of approximately 1/*N*, where *N* is sample size, so we employed an approximately unbiased estimator of LD Score given by 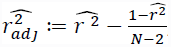, where 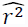 denotes the standard, biased estimator of the squared Pearson correlation^21^. Note that it is possible to have 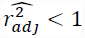, which is a mathematically necessary feature of any unbiased estimator of *r^2^.* Thus, some estimated LD Scores will be less than 1. In practice, almost all variants with estimated LD Score less than 1 were rare: only 0.01% of variants with MAF > 5% had estimated LD Scores below 1.

We examined the effect of varying the window size on our estimates of LD Score, and found that our estimates of LD Score were robust to choice of window size. The mean difference in LD Scores estimated with a 1 cM window and a 2 cM window was less than 1% of the mean LD Score (Supplementary Figure 8), and all LD Scores estimated with window sizes larger than 1 cM had squared correlations > 0.99 (Supplementary Table 7). This observation also addresses concerns about inflation in the LD Score from the intra-European population structure in the 1000 Genomes reference panel. The mean inflation in the 1 cM LD Score from population structure can be approximately bounded by the mean difference between a 1 cM LD Score and a 2 cM LD Score. Since this difference is < 1% of the mean LD Score, we conclude that bias from population structure is not significantly inflating our estimates of LD Score.

We estimated LD Score standard error via a delete-one jackknife over the 378 phased individuals in the 1000 Genomes European reference panel. We found that the LD Score standard error was positively correlated with MAF and with LD Score itself. Jackknife estimates of LD Score standard error became extremely large for variants with MAF < 1%, so we excluded variants with 1000 Genomes European sample MAF < 1% from all LD Score regressions.

### Intra-European LD Score Differences

In order to quantify the magnitude of intra-European differences in LD Score, we estimated LD Scores using each of the 1000 Genomes European subpopulations: Utah Residents with Northern and Western European Ancestry (CEU), British in England and Scotland (GBR), Toscani in Italia (TSI) and Finnish in Finland (FIN). The LD Scores from the four subpopulations were all highly correlated but the mean LD Score was not constant across populations. The mean LD Scores (MAF > 1%) were EUR, 110; CEU, 109; GBR, 104; FIN, 117; TSI, 102. The observation that the mean LD Score in the Finnish (FIN) population was elevated is consistent with a recent bottleneck in the genetic history of Finland^32^, and the observation that the mean LD Score in the Southern European TSI population is lower is consistent with reports that Southern European populations have gone through less severe bottlenecks than Northern European populations^22^.

Intra-European differences in LD Score can be a source of bias in the LD Score regression intercept. For instance, if one attempts to perform LD Score regression using the 1000 Genomes European LD Score on a GWAS with all samples from Finland, then the LD Score regression intercept may be biased upwards. Similarly, if one attempts to perform LD Score regression using the 1000 Genomes European LD Score on a GWAS with all samples from Italy, the LD Score regression intercept may be biased downwards. If we make the approximation that the intra-European differences in LD Score can be described by an additive term plus 5% noise (*i.e.,* if we assume that the FIN LD Score equals the pan-European LD Score plus seven, which is a worst-case scenario among linear relationships between the two LD Scores in terms of bias in the intercept), then the bias introduced into the LD Score regression intercept by using the pan-European LD Score to perform LD Score regression on a Finnish GWAS will be 7 multiplied by the slope of the LD Score regression plus 5% of 0.05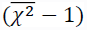 (where 7 is the difference between the reference population LD Score and the GWAS population LD Score). Since all of the mean European subpopulation LD Scores that we have estimated are within ± 8 of the mean pan-European LD Score, we estimate that the bias in the LD Score regression intercept from intra-European LD Score differences is at most ±10 times the LD Score regression slope. For the real GWAS analyzed in Table 1, this corresponds to a worst-case difference of approximately ±10% in the estimate of the proportion of the inflation in the mean *χ^2^* that results from confounding bias, with a higher probability of upward bias (because the noise term in the relationship between target and reference LD Score always causes upward bias in the LD Score regression intercept, while systematic directional differences in target and reference LD Scores can bias the LD Score regression intercept in either direction).

### LD Score Regression

We prove in the Supplementary Note that confounding bias due to population stratification and polygenic genetic architecture contribute additively to inflation in *χ^2^-*statistics. It is also true that cryptic relatedness and polygenic genetic architecture contribute additively to inflation in *χ^2^*-statistics, which we demonstrate via simulation in this paper. Therefore, we can estimate the mean contribution of confounding to test statistic inflation by regressing *χ^2^* -statistics against LD Score. This estimate will be unbiased if LD Score is uncorrelated with all sources of confounding bias, and if the assumption that the local LD Score is approximately equal in all populations from which GWAS samples were drawn. It is certainly the case that LD Score is uncorrelated with inflation due to cryptic relatedness, since inflation due to cryptic relatedness affects all SNPs equally. The inflation in the *χ*^2^-statistic of a variant from population stratification that results from a two population mixture depends on the difference in mean phenotype and the difference in allele frequency between these two populations. Under genetic drift, the expected squared difference in allele frequency between two populations (which is proportional to the inflation in *χ^2^*-statistics from stratification) is proportional to *F_ST_p*_anc_(1 − *p_anc_*),^33^ where *p_anc_* is allele frequency in the ancestral population. Note that LD Score does not appear in this expression. This quantity may be correlated with LD Score if *p_anc_* is correlated with the LD Score in the ancestral population. However, the squared correlation between *p*(1 − *p*) and LD Score in the 1000 Genomes Europeans is ~3.5%, so we expect that any correlation between LD Score and allele frequency differences due to drift will be small.

If GWAS is confounded by genetic differences between populations that result from selection, then this source of confounding will likely be correlated with LD Score, because variants with higher LD Scores are more likely to be in LD with a variant under differential selection, and are therefore more likely to show an allele frequency difference as a result of hitchhiking and because background selection causes regions with low recombination rates to drift more rapidly. However, most allele frequency differences between human population are a result of drift, with only a small contribution from selection^34,35^, so we do not expect that correlation between allele frequency differences and LD Score at sites under selection will be a major source of upward bias in the LD Score regression slope.

### Regression Weights

To account for heteroskedasticity and correlations between *χ*^2^-statistics of variants in LD, we employed a weighting scheme that gives lower weight to variants with high LD Scores.

Let *S* denote the set of variants included in the LD Score regression, and for a variant j included in *S* let 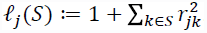. Then variant *j* is over-counted in the regression by a factor of approximately *ℓ_j_*(*S*). We estimate *ℓ_j_* (*S*) for the set of variants S described in the section *Application to Real Data* using the same procedure we used to estimate the full 1000 Genomes LD Score. Since our estimates 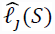 can be negative or zero, and regression weights must be positive, we weight by 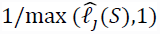 to correct for over-counting.

We observe in simulations that the variance of *χ^2^* -statistics increases with LD Score. Thus our data are heteroskedastic, and we can improve the efficiency of our regression by giving less weight to variants with high LD Score. Precisely, account we for heteroskedasticity by weighting by 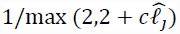, where 2 is the variance of a central *χ^2^* distribution and c is a parameter that describes the increase in variance of *χ*^2^-statistics with LD Score. We estimated *c* by binning all variants into 50 LD Score quantile bins, computing the variance of the *χ*^2^-statistics in each bin, then regressing these variances against the mean LD Score of each bin. For this paper, we take c= 0.1 for application to real GWAS and *c*=0.085 for simulations with a sample size of less than 10,000. We could obtain better regression weights by estimating *c* more accurately. The correct value of *c* depends on sample size and higher moments of the effect size distribution, and estimating these moments is the subject of ongoing research. However, we note that choice of regression weights does not change the expectations of the regression parameters, only the variance.We verified that these regression weights improve the performance of the regression by computing the standard deviation of the weighted (with c = 0.085) and unweighted regression intercepts across 100 simulations with a polygenic genetic architecture with 1% of SNPs causal and no confounding bias, so that the expected value of the LD Score regression intercept is one. The standard deviation of the unweighted regression intercept was 0.062 and the standard deviation of the weighted regression intercept was 0.015, which shows that this weighting scheme does improve the efficiency of the LD Score regression intercept as an estimator of the contribution of confounding bias to test statistic inflation.

### Attenuation Bias

Standard least-squares and weighted least-squares regression theory assumes that the explanatory variable (also referred to as the independent variable, or *X*) is measured without error. If the explanatory variable is measured with error, then the magnitude of the regression slope will be biased toward zero. This form of bias is known as attenuation bias. If the explanatory variable is measured with error, but the variance of this error is known, then it is possible to produce an unbiased regression slope by multiplying the slope by a disattenuation factor, which is equal to the squared weighted Pearson correlation between the noisy estimates of the explanatory variable and the true value of the explanatory variable. We provide an R script that can estimate this disattenuation factor given LD Scores and jackknife estimates of LD Score standard errors (see URLs).

### Regression Standard Errors

The usual estimators of regression standard errors assume uncorrelated and homoscedastic error terms, and these assumptions are violated by the LD Score regression. To produce more robust standard errors, we estimated standard errors for the weighted, disattenuated LD Score regression slope and intercept using a block jackknife over blocks of consecutive variants. We verified that our standard errors were not sensitive to the choice of block size by estimating block jackknife standard errors using block sizes between 100 and 50,000 variants and observing that the standard errors did not depend on block size. The standard errors reported in Table 1 use a delete-1000 block jackknife.

### Simulations

When performing simulations with polygenic genetic architectures using genotyped or imputed data, variants in the 1000 Genomes reference panel not included in the set of genotypes used for simulation cannot contribute to the simulated phenotypes, and so should not contribute to the LD Score used for simulations. Precisely, for the simulations with polygenicity and the simulations with polygenicity and bias, we used LD Scores where estimates of *r^2^* were derived from the 1000 Genomes European reference panel, but the sum of *r*^2’^s was taken over only those SNPs included in the simulations. For the simulations with frequency-dependent genetic architecture, we estimated LD Scores from the same genotypes used for simulations, because we wanted to quantify the bias introduced by frequency-dependent genetic architecture even when LD Scores are estimated with little noise. For the simulations with pure population stratification, we used an LD Score estimated from all 1000 Genomes variants, since there was no simulated polygenic architecture in these simulations. For simulations with pure population stratification, the details of the cohorts used are given in supplementary table 1.

It is difficult to use real genotypes to simulate ascertained studies of a binary phenotype with low population prevalence: to obtain 1000 cases with a simulated 1% phenotype, one would need to sample on expectation 100,000 genotypes, which is not feasible. We therefore generated simulated genotypes at 1.1 million SNPs with mean LD Score 110 and a simplified LD structure where *r^2^* is either 0 or 1, and all variants had 50% minor allele frequency. We generated phenotypes under the liability threshold model with all per-normalized genotype effect sizes (*i.e.,* effects on liability) drawn *i.i.d.* from a normal distribution, then sampled individuals at random from the simulated population until the desired number of cases and controls for the study had been reached. The R script that performs these simulations is available online (URLs).

### Application to Real Data

The majority of the sets of summary statistics that we analyzed did not contain information about sample minor allele frequency or imputation quality. In order to restrict to a set of common, well-imputed variants, we retained only those SNPs in the HapMap 3 reference panel^36^ for the LD Score regression. To guard against underestimation of LD Score from summing only LD with variants within a 1cM window, we removed variants in regions with exceptionally long-range LD^37^ from the LD Score regression (NB LD with these variants were included in the estimation of LD Score). Lastly, we excluded pericentromeric regions (defined as ± 3 cM from a centromere) from the LD Score regression, because these regions are enriched for sequence gaps, which may lead to underestimation of LD Score, and depleted for genes, which may reduce the probability of association to phenotype^38,39^. The final set of variants retained for LD Score regression on real data consisted of approximately 1.1 million variants.

## Acknowledgements

### AUTHOR CONTRIBUTIONS

BMN MJD NP and ALP conceived of the idea. BBS PRL SR and HF analyzed the data and performed the analyses. JY supplied code. BBS and BMN drafted the manuscript. All authors provided input and revisions for the final manuscript. MJD supplied reagents.

### COMPETING FINANCIAL INTERESTS

The authors declare no competing financial interests.

## ACKNOWLEDGEMENTS

We would like to thank P. Sullivan for helpful discussion. This work was supported by NIH grant R01 HG006399 (ALP) and US NIH grant R01 MH094421 (PGC). H. Finucane is supported by a Hertz Fellowship. Data on coronary artery disease / myocardial infarction have been contributed by CARDIoGRAMplusC4D investigators and have been downloaded from www.CARDIOGRAMPLUSC4D.ORG. Finally, we thank the coffee machine in the ATGU common area for inspiration.

## URLs

1. 1000 Genomes genetic map and haplotypes: http://mathgen.stats.ox.ac.uk/impute/data_download_1000G_phasel_integrated.html
2. LD Score database: ftp://atguftp.mgh.harvard.edu/brendan/1k_eur_r2_hm3snps_se_weights.RDS
3. Simulation and regression code: https://github.com/bulik/ld_score
4. GIANT Consortium summary statistics: http://www.broadinstitute.org/collaboration/giant/index.php/GIANT_consortium_data_files
5. PGC and TAG Consortium summary statistics: https://pgc.unc.edu/Sharing.php#SharingOpp
6. IIBDGC summary statistics (NB these summary statistics are meta-analyzed with immunochip data, which is not appropriate for LD Score regression): http://www.ibdgenetics.org/downloads.html
7. CARDIoGRAM summary statistics: http://www.cardiogramplusc4d.org/downloads/
8. DIAGRAM summary statistics: http://diagram-consortium.org/downloads.html
9. Rheumatoid Arthritis summary statistics: http://www.broadinstitute.org/ftp/pub/rheumatoid_arthritis/Stahl_etal_2010NG/
10. Blood Pressure summary statistics: http://www.georgehretlab.org/icbp_088023401234-9812599.html
11. MAGIC consortium summary statistics: http://www.magicinvestigators.org/downloads/
12. GEFOS consortium summary statistics: http://www.gefos.org/?q=content/data-release
13. SSGAC summary statistics: http://ssgac.org/Data.php

