## Supplementary Figures and Tables for "LD Score Regression Distinguishes Confounding from Polygenicity in Genome-Wide Association Studies"

*Supplementary Information*

*Table of Contents*

*Supplemental Figures	2*

*Supplementary Figure 1: Intercepts from simulations with varying heritability	2*

*Supplementary Figure 2: Slopes from simulations with varying heritability	3*

*Supplementary Figure 3: Intercepts from simulations with various proportions of causal SNPs	4*

*Supplementary Figure 4: Slopes from simulations with various proportions of causal SNPs	5*

*Supplementary Figure 5: Estimated standard error from simulations with various proportions of causal SNPs	6*

*Supplementary Figure 6: Simulations with frequency-dependent architecture	7*

*Supplementary Figure 7: Simulation where all causal variants are rare	8*

*Supplementary Figure 8: LD Score estimates with varying window size	9*

*Supplementary Tables	10*

*Supplementary Table 1: Descriptions of cohorts for simulations with pure population stratification	10*

*Supplementary Table 2: Simulations with across-cohort stratification	11*

*Supplementary Table 3: Simulations with within-cohort stratification	12*

Supplementary Table 4: Simulations with bias and polygenicity	13

*Supplementary Table 5: Simulations with Ascertained Binary Phenotypes	14*

*Supplementary Table 6: Simulations with frequency-dependent genetic architecture	15*

*Supplementary Table 7: R^2^ matrix of LD Scores with varying window sizes	16*

*Supplementary Table 8: LD Score regressions with double GC correction	17*

*Supplementary Table 9: Simulation with intergenic GC correction	18*

*Supplementary Table 10: Summary Statistic Metadata, Quantitative Trait	19*

*Supplementary Table 11: Summary Statistic Metadata, Case/Control	20*

*References*	21

### Supplemental Figures

#### Supplementary Figure 1: Intercepts from simulations with varying heritability

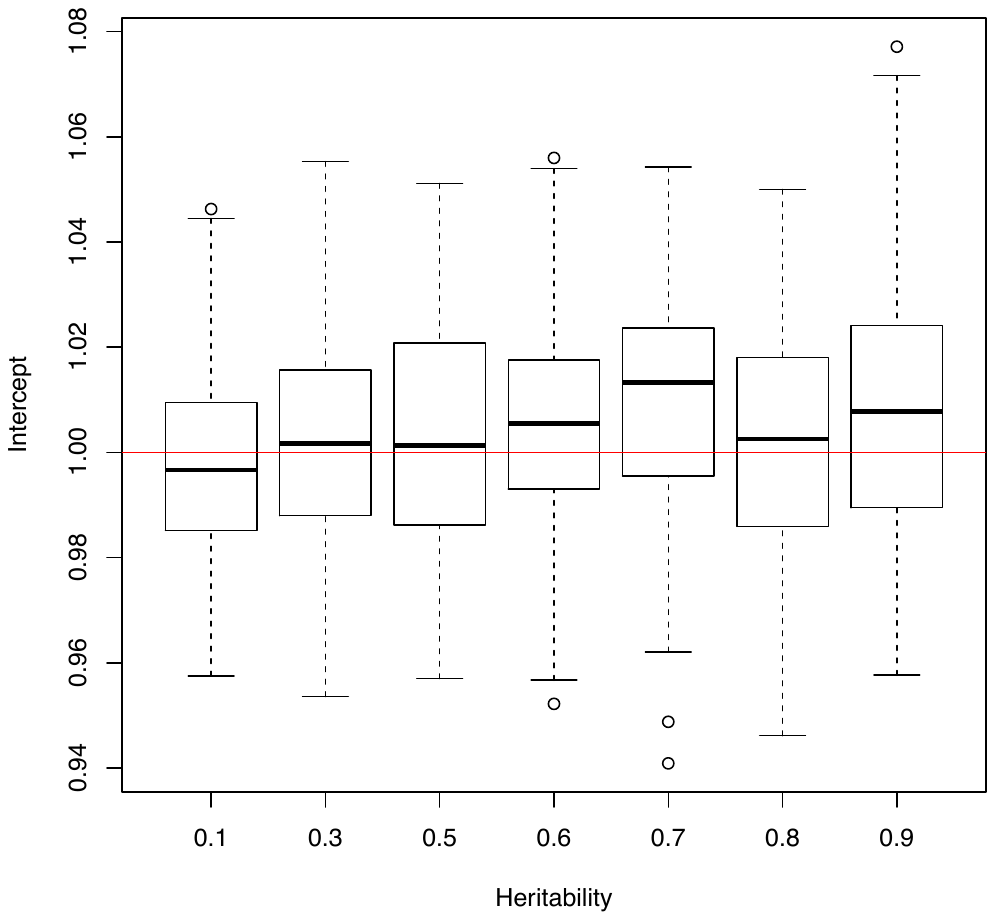
 The *x-*axis displays different heritabilities specified for simulations, and the *y*-axis displays LD Score regression intercepts from 100 simulation replicates for each value of heritability. The red line shows the expected LD Score regression intercept in the absence of confounding bias. These simulations used only SNPs on chromosome 1, which explains the large standard error. For all simulations, 1% of SNPs were causal.

#### Supplementary Figure 2: Slopes from simulations with varying heritability

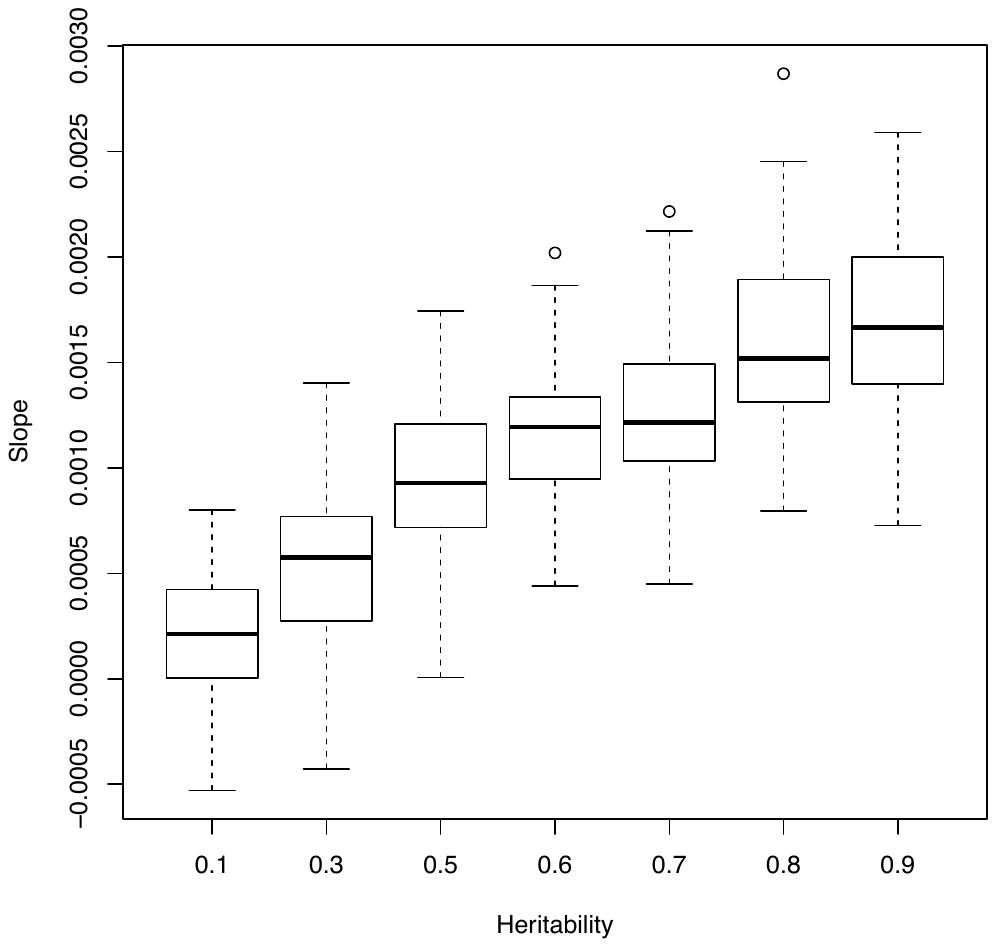

The *x-*axis displays different heritabilities specified for simulations, and the *y*-axis displays LD Score regression slopes from 100 simulation replicates for each value of heritability. These simulations used only SNPs on chromosome 1, which explains the large standard error. For all simulations, 1% of SNPs were causal.

#### Supplementary Figure 3: Intercepts from simulations with various proportions of causal SNPs

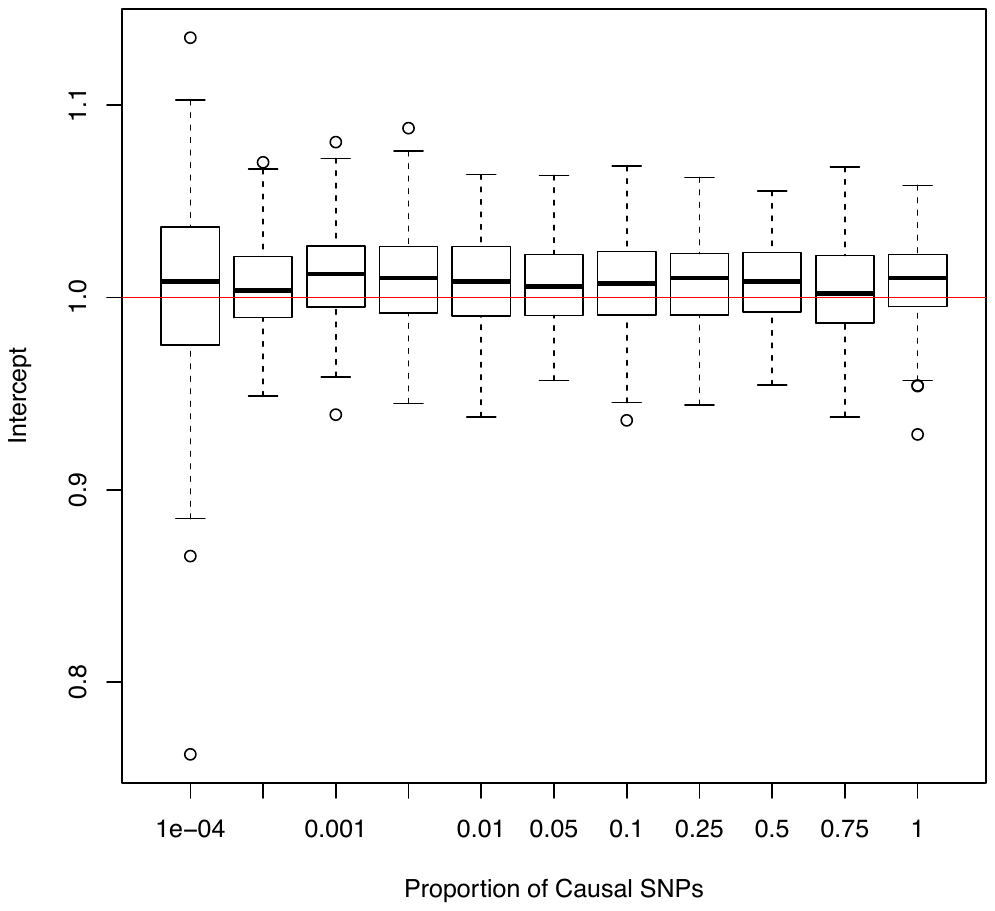
 The *x-*axis displays different proportions of causal SNPs specified for simulations, and the *y*-axis displays LD Score regression intercepts from 100 simulation replicates for each value of the proportion of causal SNPs. These simulations used only SNPs on chromosome 1, which explains the large standard error. For all simulations, the heritability was 0.9.

#### Supplementary Figure 4: Slopes from simulations with various proportions of causal SNPs

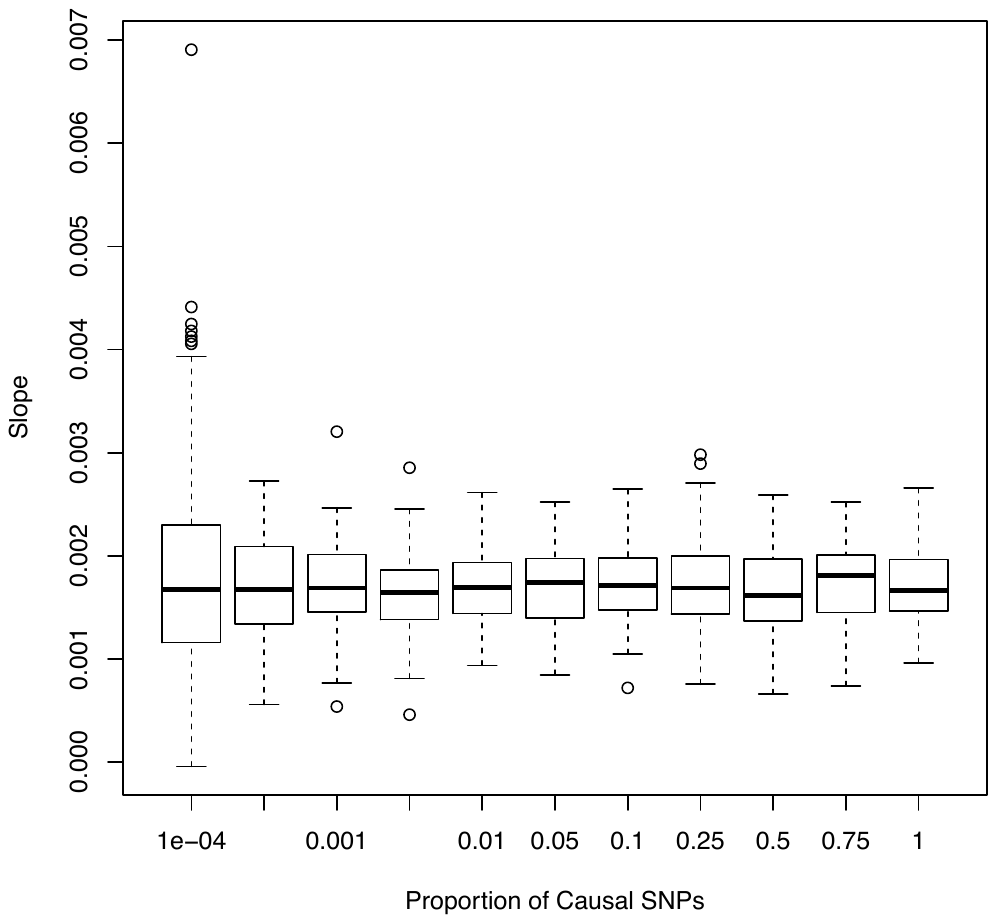

The *x-*axis displays different proportions of causal SNPs specified for simulations, and the *y*-axis displays LD Score regression slopes from 100 simulation replicates for each value of the proportion of causal SNPs. These simulations used only SNPs on chromosome 1, which explains the large standard error. For all simulations, the heritability was 0.9.

#### Supplementary Figure 5: Estimated standard error from simulations with various proportions of causal SNPs

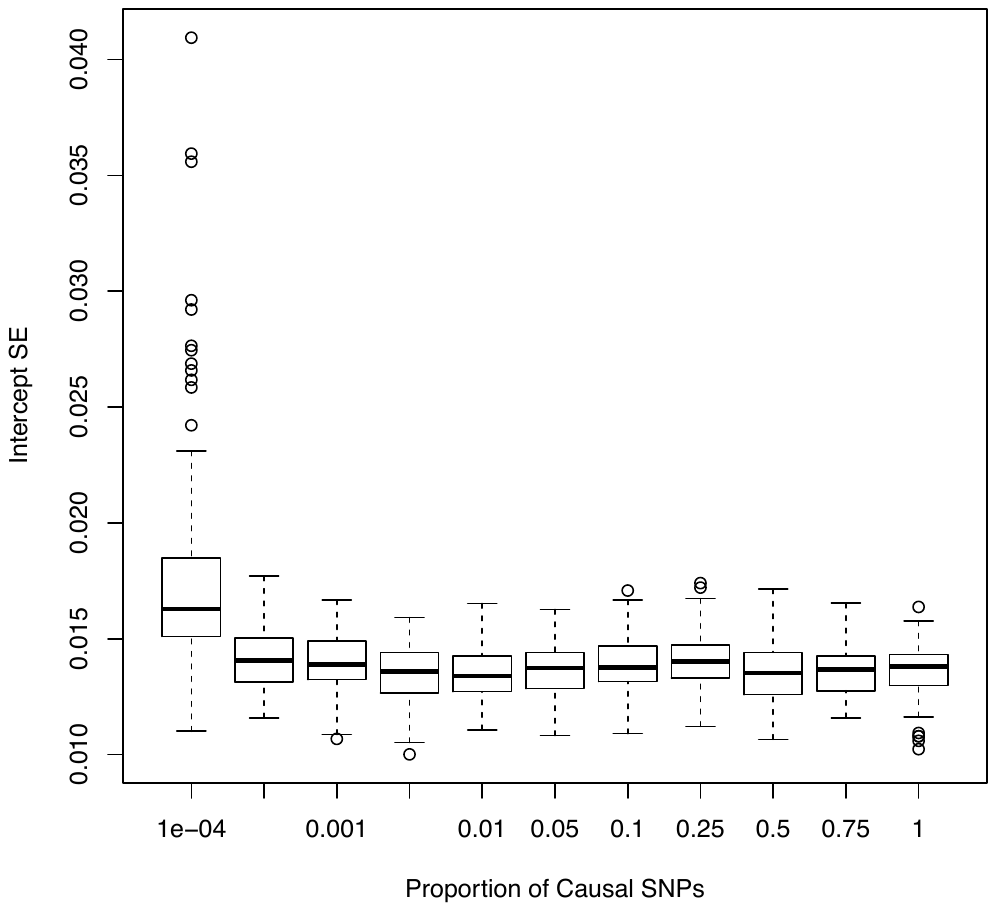

The *x-*axis displays different proportions of causal SNPs specified for simulations, and the *y*-axis displays block jackknife estimates of the standard error of the intercept from each of 100 simulation replicates for each proportion of causal SNPs. These simulations used only SNPs on chromosome 1, which explains the large standard error. For all simulations, the heritability was 0.9.

#### Supplementary Figure 6: Simulations with frequency-dependent architecture

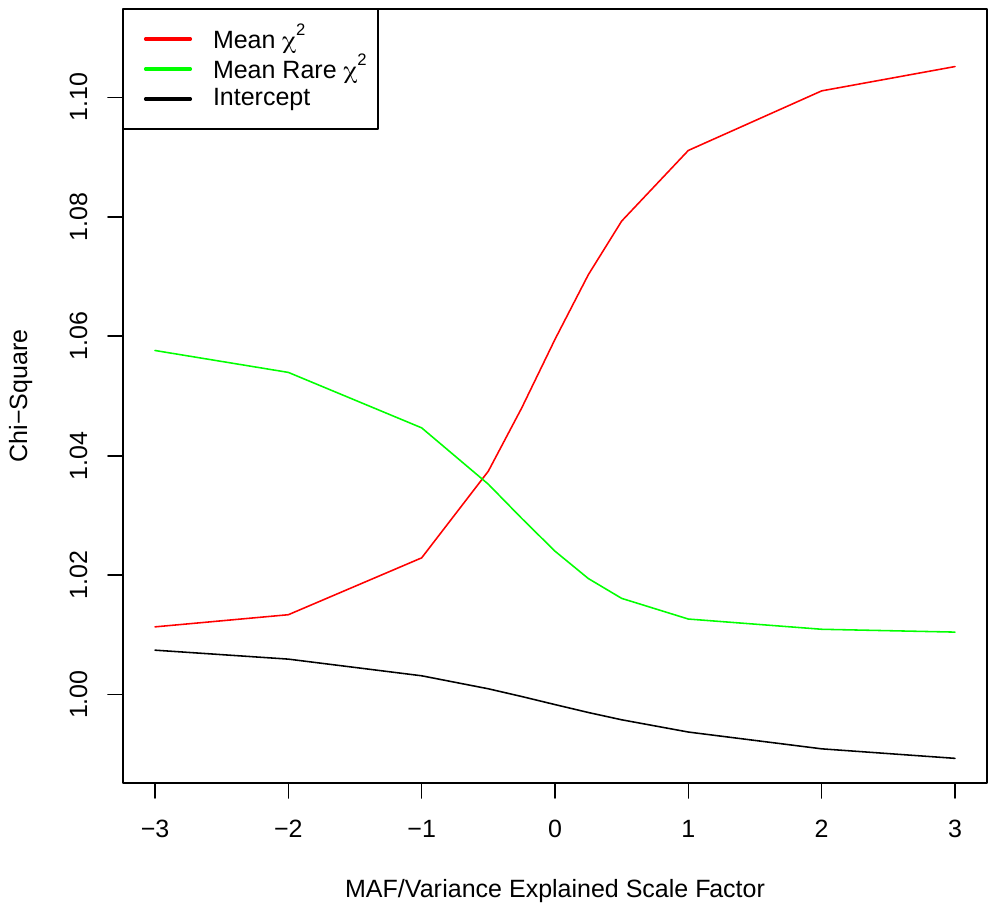

The *x*-axis describes the simulate relationship between minor allele frequency and effect size. Precisely, per-normalized genotype effects for 10,000 causal variants were drawn from$N(0,{(p(1-p))}^{x})$, where *p* is MAF and *x* is the *x*-coordinate. To prevent singleton and doubleton variants from having extreme effects for large negative values of *x*, we drew the effect sizes for variants with MAF < 1% from $N(0,0.0099^{x})$. Our model holds when *x=0.* The red line is the mean $\chi^{2}$ among the common HapMap 3^1^ variants retained for LD Score regression. The green line is the mean $\chi^{2}$ among variants with MAF < 1%. The black line is the LD Score regression intercept. Each data point is the average across 10 simulation replicates with randomly chosen causal variants and effect sizes.

#### Supplementary Figure 7: Simulation where all causal variants are rare

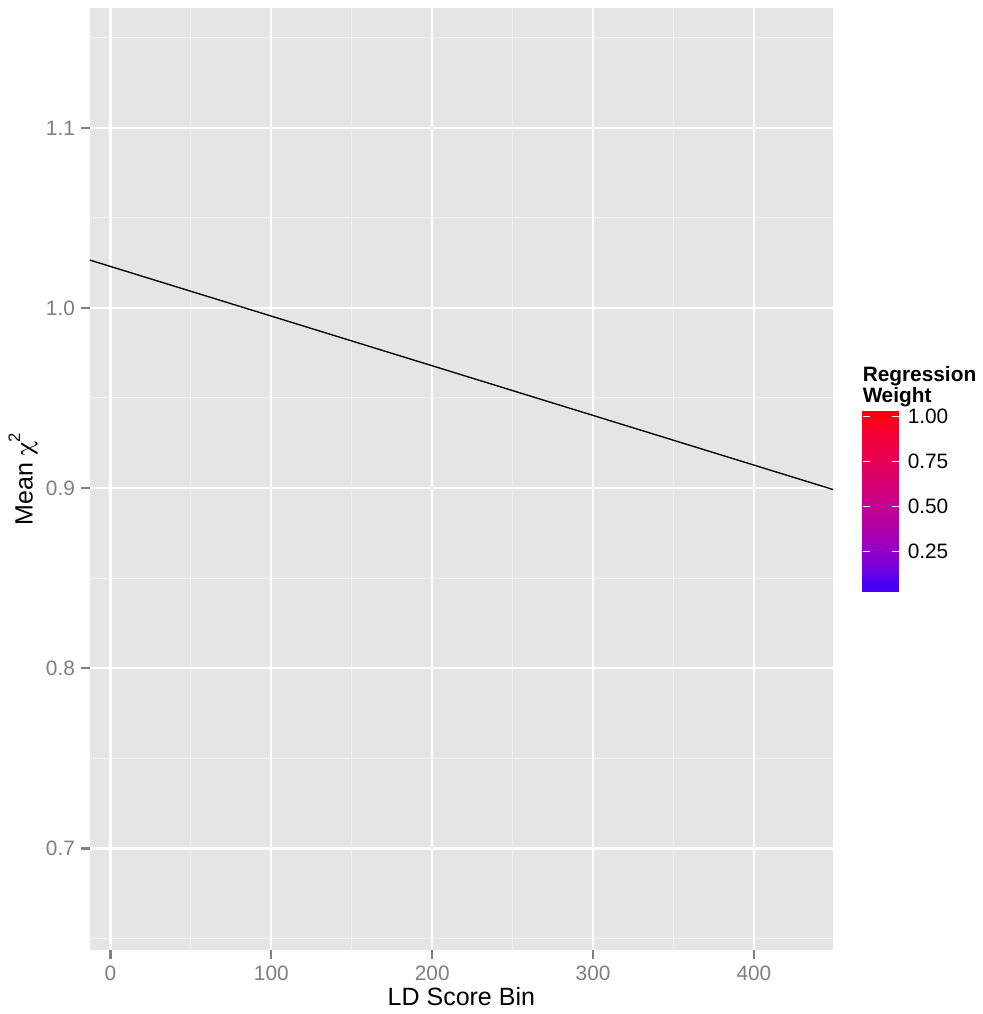

LD Score regression plot for a simulation with 1000 Swedish samples and ~700,000 SNPs on chromosome 1 where all causal variants had MAF < 1%. Each point represents an LD Score quantile, where the *x*-coordinate of the point is the mean LD Score of variants in that quantile and the *y*-coordinate is the mean $\chi^{2}$ of variants in that quantile. Colors correspond to regression weights, with red indicating large weight. The black line is the LD Score regression line. The slope of the LD Score regression line is -3.2E-4, which is statistically significantly less than zero (block jackknife *p=*0.013).

#### Supplementary Figure 8: LD Score estimates with varying window size

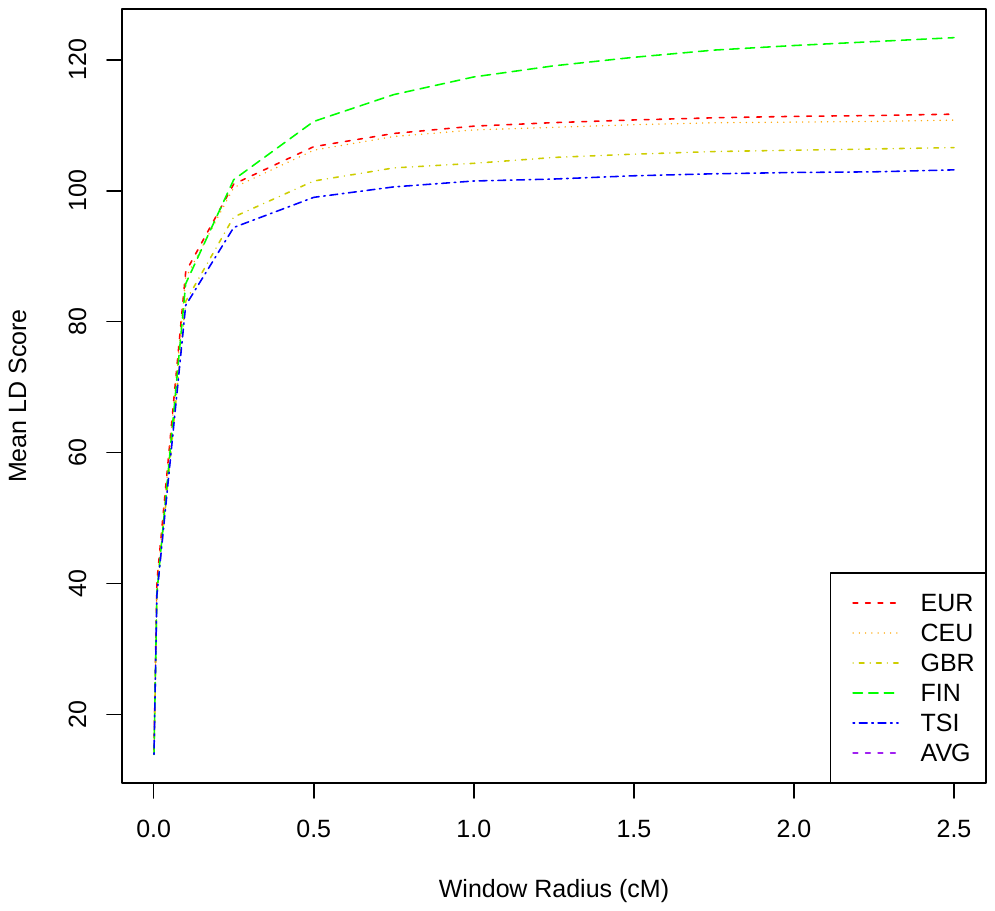

The *x-*axis displays the window radius used for estimating LD Score. The *y-*axis displays the mean LD Score among variants with sample MAF > 1% in all four 1000 Genomes European subpopulations. Each colored line represents one of the four 1000 Genomes European subpopulations: Europeans (EUR, 378 individuals), Utah Residents with Northern and Western European Ancestry (CEU, 85 individuals), British in England and Scotland (GBR, 88 individuals), Finnish in Finland (FIN, 93 individuals) and Toscani in Italia (TSI, 98 individuals). The line labeled AVG is the mean of the four subpopulation LD Scores, and is almost entirely obscured by the EUR line.

### Supplementary Tables

#### Supplementary Table 1: Descriptions of cohorts for simulations with pure population stratification

| Abbreviation | Origin | Principal Investigator | Controls |
| --- | --- | --- | --- |
| clo3 | Cardiff, UK | Walters, J | 945 |
| cou3 | UK | O’Donovan, M | 544 |
| egcu | Estonia | Esko, T | 1,177 |
| swe5 | Sweden | Sullivan, PF | 2,617 |
| swe6 | Sweden | Sullivan, PF | 1,219 |
| umeb | Umeå, Sweden | Adolfsson, R | 584 |
| umes | Umeå, Sweden | Adolfsson, R | 713 |

Supplementary table 1 describes the seven PGC Schizophrenia control cohorts used for simulation with pure population stratification. All cohorts were genotyped on the Illumina Omni Express array; only unaffected individuals (controls) and directly genotyped SNPs post-QC (between approximately 600,000 and 700,000 SNPs, depending on cohort) were retained for simulations. In total genotypes for 9,135 individuals were incorporated into the simulations with pure population stratification

#### Supplementary Table 2: Simulations with across-cohort stratification

| Population 1 | Population 2 | ${\bar{\boldsymbol{\chi}}}^{\boldsymbol{2}}$ | $\boldsymbol{(}\boldsymbol{\lambda}_{\boldsymbol{GC}}\boldsymbol{-1)/(}{\bar{\boldsymbol{\chi}}}^{\boldsymbol{2}}\boldsymbol{-1)}$ | $\boldsymbol{(\alpha-1)/(}{\bar{\boldsymbol{\chi}}}^{\boldsymbol{2}}\boldsymbol{-1)}$ |
| --- | --- | --- | --- | --- |
| cou3 | clo3 | 1.09 | 1.06 | 0.79 |
| egcu | clo3 | 9.27 | 1.02 | 0.91 |
| egcu | cou3 | 6.56 | 1.01 | 0.91 |
| swe5 | clo3 | 3.57 | 1.00 | 0.95 |
| swe5 | cou3 | 2.75 | 1.01 | 0.94 |
| swe5 | egcu | 9.36 | 1.00 | 0.93 |
| swe6 | clo3 | 3.73 | 1.00 | 0.95 |
| swe6 | cou3 | 3.84 | 0.8 | 1.14 |
| swe6 | egcu | 6.85 | 1.01 | 0.93 |
| swe6 | swe5 | 1.74 | 0.97 | 0.95 |
| umeb | clo3 | 3.72 | 0.97 | 0.96 |
| umeb | cou3 | 2.95 | 0.84 | 0.96 |
| umeb | egcu | 5.07 | 0.89 | 0.93 |
| umeb | swe5 | 2.33 | 0.86 | 0.99 |
| umeb | swe6 | 1.67 | 0.42 | 1.31 |
| umes | clo3 | 7.48 | 0.97 | 0.99 |
| umes | cou3 | 5.62 | 0.86 | 0.99 |
| umes | egcu | 10.11 | 0.91 | 0.96 |
| umes | swe5 | 7.35 | 0.93 | 1.00 |
| umes | swe6 | 4.44 | 0.92 | 1.00 |
| umes | umeb | 3.20 | 1.00 | 1.01 |
| Mean (SD) | | | **0.93 (0.l3)** | **0.98 (0.09)** |

This table describes simulations with pure across-cohort population stratification. In each simulation, individuals from population 1 were labeled cases and N­_2­_ individuals from population 2 were labeled controls. We then computed association statistics for variants in the intersection of the subset of HapMap 3 variants used for LD Score regressions on real data (Online Methods) and variants on the Illumina Omni Express array (approx. 450,000 variants in each simulation).

Column descriptions. The mean $\chi^{2}$ among these variants is displayed in the column labeled $\bar{\chi}^{2}$**.** The entries in the column labeled $(\alpha-1)/(\bar{\chi}^{2}-1)$ can be interpreted as the LD Score regression estimate of the proportion of the mean $\chi^{2}$ that results from confounding bias. This number should be close to one in simulations with pure population stratification. The entries in the column labeled $(\lambda_{GC}-1)/(\bar{\chi}^{2}-1)$ give a comparable estimate using $\lambda_{GC}$ instead of the LD Score regression intercept. The small downward bias in both $\lambda_{GC}$ and the LD Score regression intercept could result from the effects of natural selection on allele frequency differences, or from differences in phenotype definition across cohorts, if the differences are influenced by genetic factors.

#### Supplementary Table 3: Simulations with within-cohort stratification

| Population | PC | ${\bar{\boldsymbol{\chi}}}^{\boldsymbol{2}}$ | $\boldsymbol{(}\boldsymbol{\lambda}_{\boldsymbol{GC}}\boldsymbol{-1)/(}{\bar{\boldsymbol{\chi}}}^{\boldsymbol{2}}\boldsymbol{-1)}$ | $\boldsymbol{(\alpha-1)/(}{\bar{\boldsymbol{\chi}}}^{\boldsymbol{2}}\boldsymbol{-1)}$ |
| --- | --- | --- | --- | --- |
| clo3 | 1 | 2.44 | 0.69 | 1.26 |
| clo3 | 2 | 1.34 | 0.81 | 0.93 |
| clo3 | 3 | 1.31 | 0.94 | 0.95 |
| cou3 | 1 | 1.08 | 1.01 | 0.79 |
| cou3 | 2 | 1.07 | 0.90 | 0.63 |
| cou3 | 3 | 1.07 | 0.93 | 0.66 |
| egcu | 1 | 1.80 | 1.01 | 0.95 |
| egcu | 2 | 1.53 | 0.99 | 0.90 |
| egcu | 3 | 1.51 | 0.78 | 0.94 |
| swe5 | 1 | 2.80 | 0.95 | 0.94 |
| swe5 | 2 | 1.42 | 0.98 | 0.89 |
| swe5 | 3 | 1.35 | 0.94 | 0.97 |
| swe6 | 1 | 2.73 | 1.00 | 0.98 |
| swe6 | 2 | 2.51 | 0.98 | 0.95 |
| swe6 | 3 | 1.52 | 0.97 | 0.93 |
| umeb | 1 | 1.88 | 1.00 | 0.95 |
| umeb | 2 | 1.86 | 0.98 | 0.98 |
| umeb | 3 | 1.44 | 1.00 | 0.94 |
| umes | 1 | 2.03 | 1.01 | 0.93 |
| umes | 2 | 1.57 | 1.02 | 0.95 |
| umes | 3 | 1.33 | 0.98 | 0.88 |
| Mean (SD) | | | **0.95 (0.09)** | **0.92 (0.12)** |

This table describes simulations with pure within-cohort population stratification. We LD-pruned the SNPs so that no SNPs on the same chromosome had *R^2^* > 0.02, then computed the top three principal components. We then used these principal components as phenotypes and computed association statistics for the same set of variants as in the simulations described in supplementary table 2.

Column descriptions. The mean $\chi^{2}$ among these variants is displayed in the column labeled $\bar{\chi}^{2}$**.** The entries in the column labeled $(\alpha-1)/(\bar{\chi}^{2}-1)$ can be interpreted as the LD Score regression estimate of the proportion of the mean $\chi^{2}$ that results from confounding bias. This number should be close to one in simulations with pure population stratification. The entries in the column labeled $(\lambda_{GC}-1)/(\bar{\chi}^{2}-1)$ give a comparable estimate using $\lambda_{GC}$ instead of the LD Score regression intercept. The small downward bias in both $\lambda_{GC}$ and the LD Score regression intercept could result from the effects of natural selection on allele frequency differences. In addition, less-than-perfect LD pruning could account for some of the small downward bias in the LD Score regression intercept.

##### Supplementary Table 4: Simulations with bias and polygenicity

| Bias | Intercept (SD) | Null ${\bar{\boldsymbol{\chi}}}^{\boldsymbol{2}}$ (SD) | Null ${\bar{\boldsymbol{\chi}}}^{\boldsymbol{2}}$/Intercept (SD) |
| --- | --- | --- | --- |
| Relatedness | 1.46 (0.02) | 1.45 (0.02) | 1.00 (0.00) |
| Stratification | 1.53 (0.17) | 1.48 (0.15) | 0.97 (0.01) |

Column descriptions. The column labeled bias identifies the source of bias, either cryptic relatedness (from the Framingham Heart Study) or population stratification (from introducing an environmental stratification term correlated with the first PC of the WTCCC2 data). Intercept is LD Score regression intercept, with the standard deviation (SD) across five simulations in parentheses. Null $\bar{\chi}^{2}$ is the mean $\chi^{2}$ among SNPs on the opposite halves of chromosomes from causal SNPs, with SD across five simulations in parentheses. Since null SNPs are not in LD with causal SNPs, the mean $\chi^{2}$ among null SNPs precisely quantifies the mean inflation in $\chi^{2}$-statistics that results from bias. Null $\bar{\chi}^{2}$/Intercept is equal to the mean $\chi^{2}$ among null SNPs divided by the LD Score regression intercept, with the SD across five simulations in parentheses. Null $\bar{\chi}^{2}$/Intercept should be approximately equal to one if the LD Score regression intercept is accurately estimating the mean inflation in test statistics that results from bias.

#### Supplementary Table 5: Simulations with Ascertained Binary Phenotypes

| Sample Size | Prevalence | $\hat{h}_{l}^{2}$(SD) | Intercept (SD) | $\lambda_{GC}$ (SD) | $\bar{\chi}^{2}$ (SD) |
| --- | --- | --- | --- | --- | --- |
| 10000 | 0.01 | 0.804 (0.027) | 0.995 (0.048) | 2.253 (0.063) | 2.452 (0.025) |
| 10000 | 0.1 | 0.793 (0.041) | 1.006 (0.04) | 1.688 (0.031) | 1.761 (0.015) |
| 1000 | 0.01 | 0.772 (0.121) | 1.005 (0.019) | 1.139 (0.014) | 1.145 (0.015) |
| 1000 | 0.1 | 0.729 (0.226) | 1.007 (0.027) | 1.083 (0.03) | 1.076 (0.013) |

This table displays results from simulations with ascertained binary phenotypes following the liability threshold model. In all simulation replicates, the true heritability (of liability, in the population) was 0.8, the effective number of independent SNPs (defined as $M_{eff}:=M/\bar{\mathcal{l}}$) was 10,000 and the proportion of cases in the sample was 0.5. All SNPs were causal, with effect sizes (precisely, per-normalized genotype effects on liability) drawn *i.i.d.* from a normal distribution. Each entry in the table represents 20 simulation replicates. The column labeled ${\hat{\boldsymbol{h}}}_{\boldsymbol{l}}^{\boldsymbol{2}}$ lists the estimated heritability of liability in the population from the LD Score regression slope. The column labeled intercept lists LD Score regression intercepts. There was no population stratification in these simulations, so the intercept should be close to one. The columns $\boldsymbol{\lambda}_{\boldsymbol{GC}}$ and ${\bar{\boldsymbol{\chi}}}^{\boldsymbol{2}}$list the genomic control inflation factor and mean $\chi^{2}$ computed from a perfectly LD-pruned set of variants.

#### Supplementary Table 6: Simulations with frequency-dependent genetic architecture

| Exponent | Intercept (SD) | ${\bar{\boldsymbol{\chi}}}^{\boldsymbol{2}}$ (SD) |
| --- | --- | --- |
| -3 | 1.007 (0.013) | 1.011 (0.008) |
| -2 | 1.006 (0.014) | 1.013 (0.008) |
| -1 | 1.003 (0.014) | 1.023 (0.009) |
| -0.5 | 1.001 (0.013) | 1.037 (0.009) |
| -0.25 | 1.000 (0.012) | 1.048 (0.008) |
| 0 | 0.998 (0.011) | 1.059 (0.007) |
| 0.25 | 0.997 (0.011) | 1.070 (0.006) |
| 0.5 | 0.996 (0.011) | 1.079 (0.006) |
| 1 | 0.994 (0.012) | 1.091 (0.007) |
| 2 | 0.991 (0.013) | 1.101 (0.009) |
| 3 | 0.989 (0.013) | 1.105 (0.010) |

Supplementary table 5 describes simulations in which per-normalized genotype effects for 10,000 randomly chosen causal variants were drawn from$N(0,{(p(1-p))}^{x})$, where *p* is MAF and *x* is the entry in the column labeled exponent. To prevent singleton and doubleton variants from having extreme effects for large negative values of *x*, we drew the effect sizes for variants with MAF < 1% from $N(0,0.0099^{x})$. Our model holds when *x=0*. Standard errors are empirical standard errors across 10 replicates with randomly chosen causal variants and effect sizes.

#### Supplementary Table 7: *R^2^* matrix of LD Scores with varying window sizes

| cM | 0.01 | 0.1 | 0.25 | 0.5 | 0.75 | 1 | 1.25 | 1.5 | 1.75 | 2 | 2.25 | 2.5 |
| --- | --- | --- | --- | --- | --- | --- | --- | --- | --- | --- | --- | --- |
|  | 0.6677 | 0.4249 | 0.3769 | 0.3642 | 0.3543 | **0.3505** | 0.3504 | 0.3494 | 0.3489 | 0.3485 | 0.3485 | 0.3479 |
| 0.01 |  | 0.7553 | 0.6929 | 0.6732 | 0.6603 | **0.6530** | 0.6523 | 0.6502 | 0.6487 | 0.6476 | 0.6479 | 0.6463 |
| 0.1 |  |  | 0.9651 | 0.9538 | 0.9424 | **0.9359** | 0.9347 | 0.9328 | 0.9314 | 0.9303 | 0.9300 | 0.9289 |
| 0.25 |  |  |  | 0.9899 | 0.9856 | **0.9810** | 0.9801 | 0.9786 | 0.9773 | 0.9763 | 0.9764 | 0.9751 |
| 0.5 |  |  |  |  | 0.9955 | **0.9934** | 0.9930 | 0.9920 | 0.9911 | 0.9904 | 0.9904 | 0.9895 |
| 0.75 |  |  |  |  |  | **0.9981** | **0.9980** | **0.9973** | **0.9965** | **0.9960** | **0.9961** | **0.9952** |
| 1 |  |  |  |  |  |  | 0.9995 | 0.9993 | 0.9990 | 0.9986 | 0.9985 | 0.9980 |
| 1.25 |  |  |  |  |  |  |  | 0.9996 | 0.9993 | 0.9990 | 0.9990 | 0.9986 |
| 1.5 |  |  |  |  |  |  |  |  | 0.9997 | 0.9995 | 0.9994 | 0.9992 |
| 1.75 |  |  |  |  |  |  |  |  |  | 0.9997 | 0.9995 | 0.9995 |
| 2 |  |  |  |  |  |  |  |  |  |  | 0.9996 | 0.9997 |
| 2.25 |  |  |  |  |  |  |  |  |  |  |  | 0.9995 |

Each entry is the squared Pearson correlation between the LD Score estimated from the 1000 Genomes Project European reference panel with the window radii listed across the top row and the leftmost column in units of centiMorgans (cM). We chose to use the 1 cM LD Score for all LD Score regressions applied to real data, so the squared correlations with the 1 cM LD Score are in bold.

#### Supplementary Table 8: LD Score regressions with double GC correction

| Mean $\boldsymbol{\chi}^{\boldsymbol{2}}$ | $\boldsymbol{\lambda}_{\boldsymbol{GC}}$ | Intercept | Intercept SE | Slope | Slope SE | Phenotype | Ref |
| --- | --- | --- | --- | --- | --- | --- | --- |
| 1.034 | 1.014 | 0.912 | 0.00469 | 0.001185 | 5.98E-05 | Diastolic Blood Pressure | ^2^ |
| 1.037 | 1.014 | 0.899 | 0.00515 | 0.001350 | 6.40E-05 | Systolic Blood Pressure | ^2^ |
| 1.046 | 0.997 | 0.893 | 0.00498 | 0.001450 | 6.43E-05 | Femoral Neck Bone Mineral Density | ^3^ |
| 1.041 | 0.987 | 0.907 | 0.00477 | 0.001280 | 6.45E-05 | Lumbar Spine Bone Mineral Density | ^3^ |
| 1.075 | 1.009 | 0.787 | 0.00441 | 0.002756 | 6.42E-05 | Waist-Hip Ratio | ^4^ |
| 1.269 | 1.040 | 0.806 | 0.00559 | 0.004386 | 8.45E-05 | Height | ^5^ |
| 1.036 | 1.000 | 0.940 | 0.00466 | 0.000889 | 5.82E-05 | Body-Mass Index | ^6^ |
| 1.102 | 0.992 | 0.934 | 0.01162 | 0.001585 | 1.16E-04 | High-Density Lipoprotein | ^2^ |
| 1.098 | 0.990 | 0.945 | 0.01186 | 0.001535 | 1.19E-04 | Low-Density Lipoprotein | ^2^ |
| 1.021 | 0.993 | 0.942 | 0.00495 | 0.000725 | 6.25E-05 | Rheumatoid Arthritis | ^7^ |
| 1.116 | 0.991 | 0.916 | 0.00833 | 0.001995 | 9.68E-05 | Total Cholesterol | ^2^ |
| 1.116 | 0.994 | 0.891 | 0.00562 | 0.002085 | 1.02E-04 | Triglycerides | ^2^ |

This table displays LD Score regression results for studies that employed two rounds of GC correction. Unlike Table 1 in the main text, this table displays results that have no been re-inflated by the meta-analysis level GC correction factor. Entries in the column labeled $\lambda_{GC}$ may differ slightly from one, because we retained a different subset of SNPs for LD Score regression than the authors of the studies in question used to compute their GC correction factor.

#### Supplementary Table 9: Simulation with intergenic GC correction

| Annotation | Mean $\boldsymbol{\chi}^{\boldsymbol{2}}$ | Lambda |
| --- | --- | --- |
| Null (chromosome 2) | 1.0098 | 1.0082 |
| Within 100 kB of a coding exon on chromosome 1 | 1.4592 | 1.2511 |
| More than 100 kB from a coding exon on chromosome 1 | 1.2505 | **1.0817** |

This table describes a simulation with 1000 Swedish samples and ~700,000 best-guess imputed genotypes on chromosome 1. We simulated phenotypes by assigning causal effects to only SNPs within coding exons on chromosome 1. We then computed association statistics for variants within 100 kB of a gene, more than 100 kB from a gene and for null SNPs on chromosome 2. Because of long-range linkage disequilibrium, lambda (*i.e.,* $\lambda_{GC}$) is significantly elevated for intergenic SNPs even though there is no bias in the test statistics, as can be seen from the fact that the test statistics of null SNPs are not inflated.

#### Supplementary Table 10: Summary Statistic Metadata, Quantitative Trait

| Citation | Trait | N | Public | Ref |
| --- | --- | --- | --- | --- |
| Heid, *et. al.*, Nat Genet, 2010 | Waist-Hip Ratio | 113,636 | Yes | ^4^ |
| Lango Allen, *et. al.*, Nature, 2010 | Height | 183,727 | Yes | ^5^ |
| Speliotes, *et. al.*, Nat Genet, 2010 | Body Mass Index | 249,796 | Yes | ^6^ |
| TAG Consortium, Nat Genet, 2010 | Smoking | 74,053 | Yes | ^8^ |
| International Consortium for Blood Pressure GWAS, Nature, 2011 | Diastolic / Systolic Blood Pressure | 69,395 | Yes | ^2^ |
| Estrada *et. al.*, Nat Genet, 2011 | Bone Mineral Density | 32,961 | Yes | ^3^ |
| Manning *et. al.*, Nat Genet, 2012 | Fasting Insulin | 51,750 | Yes | ^9^ |
| Rietveld, *et. al.*, Science, 2013 | Years of Education | 126,559 | Yes | ^10^ |

Column descriptions. All columns are self-explanatory, except the column labeled N counts the number of individuals in the discovery phase of the GWAS, not including replication samples. The column labeled public indicates whether the summary statistics are publicly available for download (see URLs).

#### Supplementary Table 11: Summary Statistic Metadata, Case/Control

| Citation | Trait | Cases | Controls | Public | Ref |
| --- | --- | --- | --- | --- | --- |
| Neale, *et. al.*, J Am Acad Adolesc Psychiatry, 2010 | ADHD | 896 | 2455 | Yes | ^11^ |
| Stahl, *et. al.*, Nat Genet, 2010 | Rheumatoid Arthritis | 5,539 | 20,169 | Yes | ^7^ |
| PGC Bipolar Working Group, Nat Genet, 2011 | Bipolar Disorder | 4,496 | 42,422 | Yes | ^12^ |
| Schunkert *et. al.*, Nat Genet, 2011 | Coronary Artery Disease | 22,233 | 64,762 | Yes | ^13^ |
| Jostins, *et. al.*, Nature, 2012 | Inflammatory Bowel Disease | 9,968 | 20,464 | No | ^14^ |
| Jostins, *et. al.*, Nature, 2012 | Crohn’s Disease | 5,956 | 14,927 | Yes* | ^14^ |
| Jostins, *et. al.,* Nature, 2012 | Ulcerative Colitis | 6,968 | 20,464 | Yes* | ^14^ |
| Morris, *et. al.*, Nat Genet, 2012 | Type 2 Diabetes | 12,171 | 56,862 | Yes | ^15^ |
| Cross-Disorder Group, Lancet, 2013 | PGC Cross-Disorder | 33,332 | 27,888 | Yes | ^16^ |
| Ripke, *et. al.*, Mol Psych, 2013 | Major Depression | 9,240 | 9,519 | Yes | ^17^ |
| O’Donovan, *et. al.*, in preparation | Schizophrenia | 31,335** | 38,765** | No*** | ^18^ |
| Rietveld, *et. al.*, Science, 2013 | College | 22,044^****^ | 73,383 | Yes | ^10^ |

Column descriptions. All columns are self-explanatory, except the columns labeled cases and controls note the number of cases and controls in the discovery phase of the GWAS, not including replication samples. The column labeled public indicates whether the summary statistics are publicly available for download (see URLs)

* These summary statistics may be meta-analyzed with Immunochip data, which is not appropriate for LD Score regression.

** Unpublished data. These summary statistics will be made publicly available on the PGC website following publication.

*** This figure counts only European samples. The full GWAS includes several thousand Asian samples, which were excluded from the LD Score regression, because the 1000 Genomes European LD Score is not representative of LD patterns in Asian populations.

**** Here cases are individuals with college education, controls those without.
