## Supplementary Figures and Tables for "LD Score Regression Distinguishes Confounding from Polygenicity in Genome-Wide Association Studies"

### SUPPLEMENTARY NOTE

#### 1. LD Score in an Unstructured Sample

1.1. **Model.** We model phenotypes as generated from the equation

$$\phi = X\beta + \epsilon, \tag{1.1}$$

where $\phi$ is an $N \times 1$ vector of (quantitative) phenotypes, $X$ is an $N \times M$ matrix of genotypes normalized to mean zero and variance one (we ignore the distinction between normalizing and centering in our sample and in the population since the error so introduced has expectation zero and $\mathscr{O}(1/N)$ variance), $\beta$ is an $M \times 1$ vector of per-normalized-genotype effect sizes and $\epsilon$ is an $N \times 1$ vector of environmental (or non-genotyped genetic) effects. We describe a model where all three variables on the right side of equation 1.1 are random. In this model, $\mathbb{E}[\epsilon] = 0$ , $\mathrm{Var}[\epsilon] = (1-h_g^2)I$, $\mathbb{E}[\beta] = 0$ and $\mathrm{Var}[\beta] = (h_g^2/M)I$[1]. To model genotypes, we assume that the genotype at variant $j$ for individual $i$ is independent of other individuals' genotypes, but we do incorporate linkage disequilibrium into the model: define $r_{jk} := \mathbb{E}[X_{ij}X_{ik}]$, which does not depend on $i$. Finally, we assume that $X$, $\beta$ and $\epsilon$ are mutually independent. We will relax the assumption that environmental effects are independent of genotype when we model population stratification in §2.2.

1.2. **Relationship between LD and $\chi^2$-Statistics.** For each variant $j = 1, \dots, M$, we compute least-squares estimates of effect size $\hat{\beta}_j := X_j^\intercal \phi / N$ (where $X_j$ denotes the $N \times 1$ vector of genotypes at variant $j$) and $\chi^2$-statistics $\chi_j^2 := N\hat{\beta}_j^2$. In this section, we compute $\mathbb{E}[\chi_j^2]$ with the expectation taken over random $X$, $\beta$, $\epsilon$.

Since $\mathbb{E}[\hat{\beta}_j] = 0$, observe that $\mathbb{E}[\chi_j^2] = N \cdot \mathrm{Var}[\hat{\beta}_j]$. We will obtain the variance of $\hat{\beta}_j$ via the law of total variance:

$$\begin{aligned} \mathrm{Var}[\hat{\beta}_j] &= \mathbb{E}[\mathrm{Var}[\hat{\beta}_j \mid X]] + \mathrm{Var}[\mathbb{E}[\hat{\beta}_j \mid X]] \\ &= \mathbb{E}[\mathrm{Var}[\hat{\beta}_j \mid X]], \end{aligned} \tag{1.2}$$

where the second line follows from the fact that $\mathbb{E}[\hat{\beta}_j \mid X] = 0$ irrespective of $X$. First,

$$\begin{aligned} \mathrm{Var}[\hat{\beta}_j \mid X] &= \frac{1}{N^2}\mathrm{Var}[X_j^\intercal \phi \mid X] \\ &= \frac{1}{N^2} X_j^\intercal \mathrm{Var}[\phi \mid X] X_j \\ &= \frac{1}{N^2}\left(\frac{h_g^2}{M} X_j^\intercal X X^\intercal X_j + N(1-h_g^2)\right). \end{aligned} \tag{1.3}$$

---

[1] This is the same assumption made in [1]. If one wishes to specify a different variance structure for the per-normalized-genotype effect sizes, *e.g.*, $\mathrm{Var}[\beta_j] = f_j$, then all results presented herein hold with normalized genotypes $(G_{ij} - 2p_j)/\sqrt{f_j}$ replacing the usual $(G_{ij} - 2p_j)/\sqrt{2p_j(1-p_j)}$, where $G_{ij}$ denotes additively coded (0,1,2) genotypes.

We can write the term on the left in terms of more familiar quantities as

$$\frac{1}{N^2} X_j^{\mathsf{T}} X X^{\mathsf{T}} X_j = \sum_{k=1}^{M} \tilde{r}_{jk}^2, \tag{1.4}$$

where $\tilde{r}_{jk} := \frac{1}{N}\sum_{i=1}^{N} X_{ij} X_{ik}$ denotes the sample correlation between variants $j$ and $k$. Define the LD Score of variant $j$ as

$$\ell_j := \sum_{k=1}^{M} r_{jk}^2. \tag{1.5}$$

Since

$$\mathbb{E}[\tilde{r}_{jk}^2] \approx r_{jk}^2 + (1 - r_{jk}^2)/N \tag{1.6}$$

(where the approximation sign hides terms of order $\mathscr{O}(1/N^2)$ and smaller; one can obtain this approximation via *e.g.,* the $\delta$-method),

$$\mathbb{E}\left[\sum_{k=1}^{M} \tilde{r}_{jk}^2\right] \approx \ell_j + \frac{M - \ell_j}{N}. \tag{1.7}$$

Thus,

$$\begin{aligned} \mathbb{E}[\chi_j^2] &\approx \frac{N(1-1/N)h_g^2}{M}\ell_j + 1 \\ &\approx \frac{N h_g^2}{M}\ell_j + 1, \end{aligned} \tag{1.8}$$

Values of $N$ considered in the main text generally fall between $10^4$ and $10^5$, so the approximation $1 - 1/N \approx 1$ is appropriate.

### 2. LD Score with Population Stratification

2.1. **Model of Population Structure.** We model population structure induced by genetic drift in a mixture of two populations in equal proportions. We draw a matrix of normalized genotypes $X$ consisting of $N/2$ samples from population 1 and $N/2$ samples from population 2 (we will use the notation $i \in P_m$ for $m \in \{1, 2\}$ to denote that individual $i$ is a member of population $m$), subject to the following constraints: $\text{Var}[X_{ij}] = 1$, $\mathbb{E}[X_{ij} \mid i \in P_1] = f_j$ and $\mathbb{E}[X_{ij} \mid i \in P_2] = -f_j$. We model the drift term $f$ as $f \sim N(0, F_{ST}V)$, where $V$ is a correlation matrix and $F_{ST}$ is Wright's $F_{ST}$ [2]. We postpone discussion of the off-diagonal entries of $V$ (which might depend on LD in the ancestral population or recombination rates) until §2.2. Finally, if $\ell_{j,m}$ denotes the LD Score of variant $j$ in population $m$, we assume that $\ell_{j,1} \approx \ell_{j,2} =: \ell_j$. The last assumption warrants a brief explanation. Assuming approximately equal LD Scores in both populations is certainly not reasonable for very large values of $F_{ST}$ (*e.g.,* if population 1 and population 2 are from different continents) or in scenarios where one population has passed through a more severe bottleneck than the other (*e.g.,* if population 1 is from Finland and population 2 is from West Africa). However, we are interested in modeling the population stratification that may remain after principal components analysis[2] in GWAS that sample from non-admixed populations, and for this purpose the assumption that $\ell_{j,1} \approx \ell_{j,2}$ seems reasonable,

[2]Including principal components as covariates in GWAS is equivalent to regressing PC-residualized phenotypes against PC-residualized genotypes. PC-residualized genotypes will have more homogeneous LD structure than the raw genotypes.

and is supported by the large values of $R^2(\ell_{j,m}, \ell_{j,n})$ that we observe for all pairs $(m, n)$ of 1000 Genomes European subpopulations.

For reference, typical values of $F_{ST}$ for human populations are $\approx 0.1$ for populations from different continents [2], $\approx 0.01$ for populations on the same continent [3], and $< 0.01$ for subpopulations within the same country [4].

2.2. **LD in a Mixture of Populations.** Suppose $j$ and $k$ are unlinked variants such that $r_{jk,1} = r_{jk,2} = 0$ and $f_j$ is independent of $f_k$. In a mixture of populations, it will often hold that $j$ and $k$ will be in LD in the whole population even if they are in equilibrium in both component populations. Given fixed $f$,

$$\begin{aligned} \mathbb{E}[r_{jk} \mid f] &= \mathbb{E}[X_{ij}X_{ik} \mid f] \\ &= \frac{1}{2}\left(\mathbb{E}[X_{ij}X_{ik} \mid f, i \in P_1] + \mathbb{E}[X_{ij}X_{ik} \mid f, i \in P_2]\right) \\ &= f_j f_k. \end{aligned} \tag{2.1}$$

If we take the expectation over random $f_j$ and $f_j$, then $\mathbb{E}[r_{jk}] = 0$, because $f_j$ and $f_k$ are independent with expectation zero. We can use equation 2.1 to compute the variance,

$$\begin{aligned} \mathrm{Var}[r_{jk}] &= \mathrm{Var}[\mathbb{E}[r_{jk} \mid f]] + \mathbb{E}[\mathrm{Var}[r_{jk} \mid f]] \\ &= \mathbb{E}[f_j^2 f_k^2] + 0 \\ &= \mathbb{E}[f_j^2]\mathbb{E}[f_k^2] \\ &= F_{ST}^2. \end{aligned} \tag{2.2}$$

Observe that since $\mathbb{E}[r_{jk}] = 0$, $\mathrm{Var}[r_{jk}] = \mathbb{E}[r_{jk}^2]$. By equation 1.5, in a finite sample,

$$\mathbb{E}[\tilde{r}_{jk}^2] \approx F_{ST}^2 + (1 - F_{ST}^2)/N. \tag{2.3}$$

Thus the sample LD Score is approximately

$$\begin{aligned} \mathbb{E}[\tilde{\ell}_j] &\approx \ell_j + MF_{ST}^2 + \frac{M(1 - F_{ST}^2)}{N} \\ &\approx \ell_j + MF_{ST}^2 + \frac{M}{N}. \end{aligned} \tag{2.4}$$

Note that we have ignored the case where $j$ and $k$ are linked and $V_{jk} \neq 0$. In this case, $\mathbb{E}[f_j^2 f_k^2] = F_{ST}^2 + 2F_{ST}V_{jk}^2$ (from the formula for the double second moments of a multivariate normal distribution). Even if for some variants $j$, the number of variants $k$ such that $V_{jk} > 0$ is $\approx 10^3$, this will make a negligible difference in $\mathbb{E}[\tilde{\ell}_j]$, because $\sum_{k: V_{jk} > 0} 2F_{ST}V_{jk}^2 < 2000F_{ST} \ll MF_{ST}$ when $M \approx 10^7$.

2.3. **Model of Stratified Phenotype.** We model phenotypes as generated by the equation

$$\phi = X\beta + S + \epsilon, \tag{2.5}$$

where $X$ is as described in §2.1, $\beta$ is as described in §1.1 and where we introduce an environmental stratification term $S$, defined by

$$S_i := \begin{cases} \sigma_s/2 & i. \in P_1 \\ -\sigma_s/2, & i \in P_2. \end{cases} \tag{2.6}$$

Finally, $\epsilon$ is as described in §1.1, except $\mathrm{Var}[\epsilon] = (1 - h_g^2 - \sigma_s^2)$, which assures that the variance of $\phi$ in the population is 1. Note that we have implicitly required that parameters be chosen such that $1 - h_g^2 - \sigma_s^2 \geq 0$.

2.4. **Relationship between LD and Stratified $\chi^2$-Statistics.** We compute $\chi^2$-statistics as defined in §1.1. In this section, we compute $\mathbb{E}[\chi_j^2]$ with the expectation taken over random $X$, $\beta$, $\epsilon$, $f$ but with $S$ fixed to ensure population stratification. Since $\mathbb{E}[\hat{\beta}_j] = 0$, observe that $\mathbb{E}[\chi_j^2] = N \cdot \mathrm{Var}[\hat{\beta}_j]$. We will obtain the variance of $\hat{\beta}_j$ via the law of total variance:

$$\mathrm{Var}[\hat{\beta}_j] = \mathbb{E}[\mathrm{Var}[\hat{\beta}_j \,|\, X]] + \mathrm{Var}[\mathbb{E}[\hat{\beta}_j \,|\, X]]. \tag{2.7}$$

Note that one can calculate $f$ from $X$, so by conditioning on $X$ we also implicitly condition on $f$. Unlike in equation 1.2, $\mathbb{E}[\hat{\beta}_j \,|\, X] \neq 0$, because of confounding from population stratification. The inner portion of the first term on the right side of equation 2.6 is the same as in equation 1.3,

$$\mathrm{Var}[\hat{\beta}_j \,|\, X] = \frac{1}{N^2}\left(\frac{h_g^2}{M} X_j^\top X X^\top X_j + N(1 - h_g^2)\right). \tag{2.8}$$

We can take the expectation over random $X$, $p$, $q$, using the result from §2.2 that in a sample from a two-way mixture of populations,

$$\frac{1}{N^2}\mathbb{E}[X_j^\top X X^\top X_j] \approx \ell_j + M F_{ST}^2 + \frac{M}{N} \tag{2.9}$$

Thus,

$$\begin{aligned} \mathbb{E}[\mathrm{Var}[\hat{\beta}_j \,|\, X]] &= \frac{1}{N^2}\left(\frac{h_g^2}{M}\mathbb{E}[X_j^\top X X^\top X_j] + N(1 - h_g^2)\right) \\ &\approx \frac{h_g^2}{M}\ell_j + h_g^2 F_{ST}^2 + \frac{1}{N} \end{aligned} \tag{2.10}$$

Next, the inner portion of the second term on the right side of equation 2.6 is

$$\begin{aligned} \mathbb{E}[\hat{\beta}_j \,|\, X] &= \frac{1}{N}\mathbb{E}[X_j^\top X\beta + X_j^\top S + X_j^\top \epsilon] \\ &= \frac{1}{N} X_j^\top S \\ &= f\sigma_s. \end{aligned} \tag{2.11}$$

Since $f$ has variance $F_{ST}$, $\mathrm{Var}[f\sigma_s] = \sigma_s^2 F_{ST}$. Thus,

$$\begin{aligned} \mathbb{E}[\chi_j^2] &= N \cdot \mathrm{Var}[\hat{\beta}_j] \\ & \frac{N h_g^2}{M}\ell_j + 1 + N F_{ST}(\sigma_s^2 + h_g^2 F_{ST}). \end{aligned} \tag{2.12}$$

We can interpret the final term, $N F_{ST}(\sigma_S^2 + h_g^2 F_{ST})$, as $N F_{ST}$ times the expected squared mean difference in phenotype between populations, which has environmental component $\sigma_s^2$ and genetic component $h_g^2 F_{ST}$ (if we model $X$, $\beta$ and $f$ as random, there is zero genetic stratification on expectation, but with some small variance about zero). Precisely, if we let $\bar{\phi}_m$ denote the mean phenotype in population $m \in \{1,2\}$, then

$$\mathbb{E}[(\bar{\phi}_1 - \bar{\phi}_2)] = \sigma_s^2 + \sum_{j=1}^{M} \mathbb{E}[\beta_j^2] \left( \sum_{i \in P_1} \mathbb{E}[X_{ij}^2 \mid i \in P_1] + \sum_{i \in P_2} \mathbb{E}[X_{ij}^2 \mid i \in P_2] \right) \tag{2.13}$$
$$= \sigma_s^2 + h_g^2 F_{ST}.$$

Set $a := \mathbb{E}[(\bar{\phi}_1 - \bar{\phi}_2)^2]$. Then we have

$$\mathbb{E}[\chi_j^2] = \frac{N h_g^2}{M} \ell_j + 1 + aNF_{ST}. \tag{2.14}$$
